# GEMC1-MCIDAS transcriptional program regulates multiciliogenesis in the choroid plexus and acts as a barrier to tumorigenesis

**DOI:** 10.1101/2020.11.22.393298

**Authors:** Qun Li, Zhiyuan Han, Navleen Singh, Berta Terré, Ryann M. Fame, Uzayr Arif, Thomas D. Page, Tasneem Zahran, Ahmed Abdeltawab, Yuan Huang, Ping Cao, Jun Wang, Hao Lu, Hart G.W. Lidov, Kameswaran Surendran, Lizhao Wu, Ulrich Schüller, Robert J. Wechsler-Reya, Maria K. Lehtinen, Sudipto Roy, Zhongmin Liu, Travis H. Stracker, Haotian Zhao

## Abstract

Multiciliated cells (MCCs) in the brain include the ependymal cells and choroid plexus (CP) epithelial cells. The CP secretes cerebrospinal fluid that circulates within the ventricular system, driven by ependymal cilia movement. However, the mechanisms and functional significance of multiciliogenesis in the CP remain unknown. Deregulated oncogenic signals cause CP carcinoma (CPC), a rare but aggressive pediatric brain cancer. Here we show that aberrant NOTCH and Sonic Hedgehog signaling in mice drive tumors that resemble CPC in humans. NOTCH-driven CP tumors were monociliated, whereas disruption of the NOTCH complex restored multiciliation and decreased tumor growth. NOTCH suppressed multiciliation in tumor cells by inhibiting the expression of GEMC1 and MCIDAS, early regulators of multiciliogenesis. Consistently, GEMC1-MCIDAS function is essential for multiciliation in the CP, and is critical for correcting multiciliation defect in tumor cells by a NOTCH inhibitor. Disturbances to the *GEMC1* program are commonly observed in human CPCs characterized by solitary cilia. Consistently, CPC driven by deletion of *Trp53* and *Rb1* in mice exhibits a cilia deficit consequent to loss of *Gemc1-Mcidas* expression. Taken together, these findings reveal a GEMC1-MCIDAS multiciliation program in the CP critical for inhibiting tumorigenesis, and it may have therapeutic implications for the treatment of CPC.

## Introduction

MCCs on the epithelial lining of the brain ventricles, the airway, and reproductive tracts control fluid flow through the synchronized beating of multiple motile cilia on their apical surface. Formation of these multiple cilia involves massive production of centrioles that subsequently migrate and dock with the apical membrane, where they nucleate ciliary differentiation. Multiciliogenesis is directed by a hierarchical network of transcriptional and post-transcriptional regulators that collaborate to exert precise control of the synthesis, directional transport and assembly of a diverse set of structural and functional components of cilia (Spassky and Meunier 2017; Lewis and Stracker 2020). Geminin Coiled-Coil Domain Containing 1 (GEMC1) and multi-ciliate differentiation and DNA synthesis associated cell cycle protein (MCIDAS or Multicilin), members of the Geminin family of coiled-coil containing nuclear proteins, are crucial early transcriptional regulators of the MCC fate (Kyrousi et al. 2015; Zhou et al. 2015; Arbi et al. 2016; Terre et al. 2016; Lu et al. 2019). GEMC1 and MCIDAS, complexed with cognate transcription partner E2Fs (E2F4 and E2F5), activate a network of downstream MCC factors including forkhead box J1 (FOXJ1), v-myb avian myeloblastosis viral oncogene homolog (MYB), and cyclin O (CCNO) (Tan et al. 2013; Boon et al. 2014; Ma et al. 2014; Wallmeier et al. 2014). Recent studies also showed that *TAp73*, a P53 family member that acts downstream of *Gemc1* in multiciliogenesis, plays an important role in the MCC program through activation of downstream genes including *Foxj1* and *Myb* (Jackson and Attardi 2016; Marshall et al. 2016; Nemajerova et al. 2016; Lalioti et al. 2019a). Disruption of the GEMC1-MCIDAS network leads to defective multiciliogenesis in different tissues, whereas mutations in both *GMNC*, the human ortholog of *GEMC1*, and *MCIDAS* have been identified in human ciliopathies (Spassky and Meunier 2017; Lewis and Stracker 2020).

In the brain, GEMC1-MCIDAS network regulates multiciliogenesis in ependymal cells (Kyrousi et al. 2015; Kyrousi et al. 2016; Terre et al. 2016), whereas Geminin and GEMC1 play antagonistic roles to maintain neural stem cells and ependymal cells in the adult neurogenic niche (Lalioti et al. 2019b; Ortiz-Alvarez et al. 2019). Interestingly, MCCs are present in the CP, a highly vascularized epithelial tissue in vertebrate brain ventricles (Narita and Takeda 2015). The CP is responsible for the synthesis and secretion of cerebrospinal fluid (CSF) that protects the internal environment of the central nervous system, whereas the movement of multiple motile cilia on the ependymal cells drives CSF flow within the ventricular system. The CSF has been increasingly recognized as an important source of molecules and morphogens that influences self-renewal and differentiation decisions of adult neural stem cells (Fame and Lehtinen 2020). Homeostatic regulation of the CSF, glial and neuronal cells plays a pivotal role in normal and abnormal brain function. Dysfunction of CP barrier properties is thought to contribute to a variety of neurological conditions (Liddelow 2015). Studies with the mouse have revealed that unlike motile cilia of ependymal cells, the characteristic multiple cilia clusters in the CP are not involved in motile functions to control CSF flow (Nonami et al. 2013). Indeed, the CP MCCs are quite unlike MCCs described in other tissues: they arise around embryonic (E) day 15, display increased motility of their multiple cilia until birth, and experience a gradual regression in the motility during post-natal life. Ultrastructural analysis has revealed that these cilia have a mixture of (9+2), (9+0) as well as (9+1) configuration of axonemal microtubules when they exhibit motility (this ultrastructural variation apparent even within a single MCC), and then switch to the (9+0) arrangement in the majority of the cilia as they lose motility (Narita et al. 2012; Nonami et al. 2013). Moreover, TAp73 is expressed in early MCCs in the CP as roof plate progenitors exit the cell cycle and initiate multiciliogenesis. However, *TAp73* is dispensable for MCC differentiation in the CP (Wildung et al. 2019). Therefore, the molecular mechanisms governing multiciliogenesis in the CP and functional significance of these cilia in health and diseases remain poorly understood.

Like other epithelial cells, the CP is susceptible to oncogenic processes. Tumors of the CP are rare primary brain neoplasms mostly found in children, but can occur at any age. CP tumors comprise up to 20% of brain tumors diagnosed in children under one year of age. Though CP papilloma (CPP) is more benign, CP carcinoma (CPC) is an aggressive tumor with a poor survival rate and a tendency for recurrence and metastasis (Gozali et al. 2012; Zaky and Finlay 2018). Research aimed at mechanism-based therapies for CPC addresses the pressing need for improved outcome. A subset of CP tumors exhibit abnormal NOTCH pathway activity (Beschorner et al. 2013). Using animal models, we have previously demonstrated that sustained NOTCH1 expression led to CPP arising from monociliated progenitors in the roof plate that proliferate in response to Sonic Hedgehog (SHH) (Eberhart 2016; Li et al. 2016). Here, we show that persistent NOTCH and SHH signals in mice drive aggressive tumors that are similar to CPC in humans (Zhu et al. 2017; Taher et al. 2019). These CP tumors display singular primary cilia resulting from repression of the GEMC1-MCIDAS transcriptional network by NOTCH, whereas biochemical or pharmacological disruption of the NOTCH complex, or *Gemc1-Mcidas* overexpression, restored multiciliation and suppressed tumor cell proliferation. Consistent with these findings, GEMC1-MCIDAS molecular network was essential for MCC differentiation in the CP, and *Gemc1* deficiency impaired the restoration of the multiciliation defect in tumor cells by a NOTCH inhibitor. Likewise, *Trp53*-deficient CPC in mice and CPC in humans exhibit identical cilia defects and a deficient GEMC1-directed transcriptional program. These findings underscore a critical role of a compromised GEMC1-MCIDAS multiciliogenesis program in CPC, and suggest that this could be exploited therapeutically to impair tumor proliferation and promote tumor differentiation.

## Results

### Gemc1-Mcidas signaling cascade is essential for multiciliated epithelia in the CP

To determine the role of *Gemc1* in the differentiation of MCCs in the CP, we examined the brains of *Gemc1*-null mice. Consistent with previous studies, transmission electron microscopy and immunostaining revealed singular cilia in ependymal cells in the walls lining the brain ventricles of *Gemc1*-null animals, whereas ependymal cells in wild type animals displayed multiple motile cilia (Supplemental Fig. S1A,B). Instead of multiciliated epithelial cells, CP in *Gemc1*-deficient animals was comprised of monociliated cells (Fig. 1A,B; Supplemental Fig. S1B; Supplemental Fig. S2A,B). The expression of CP lineage markers including orthodenticle homeobox 2 (OTX2), aquaporin 1 (AQP1), transthyretin (TTR) and cytokeratins was comparable between *Gemc1*^−/−^ and wild type CP (Fig. 1C-E; Supplemental Fig. S2C). In wild type animals, *Gemc1* mRNA was detected in epithelial cells adjoining the roof plate. However, *Gemc1* expression persisted in the CP epithelium and in ependymal cells of *Gemc1*-deficient animals, likely due to expression of mutant *Gemc1* transcripts that may be upregulated by a compensatory pathway (Fig. 2A; Supplemental Fig. S2D; Supplemental Fig. S3A). Nonetheless, the expression of *Foxj1* and TAp73 in multiciliated epithelial and ependymal cells was significantly reduced in the absence of *Gemc1*, whereas *Gemc1* overexpression stimulated TAp73 expression (Fig. 2A,B; Supplemental Fig. S2D-F; Supplemental Fig. S3B,D). Thus, *Gemc1* expression in the CP epithelium critically mediates the expression of *TAp73* and *Foxj1*, and is essential for the differentiation of MCCs in the CP.

**Figure 1.**
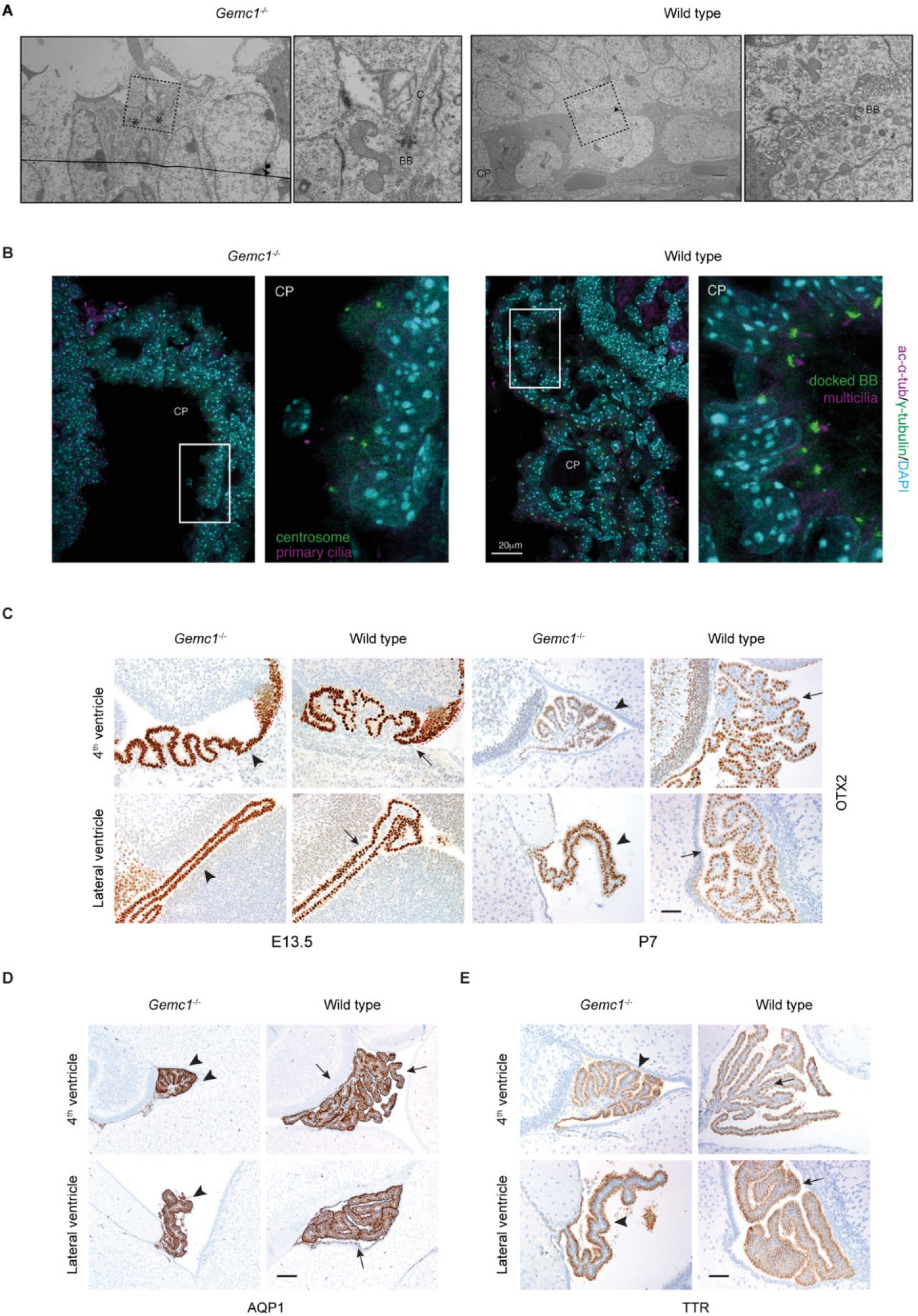
Loss of multiciliated cells in *Gemc1*-null CP. (*A*) Transmission electron micrographs are shown of CP epithelial cells in newborn *Gemc1*^*−/−*^ (asterisks) and wild type (arrow) mice. Boxed regions are magnified on the right. C, cilia; BB, basal body. (*B*) The expression of acetylated α-tubulin (ac-α-tub, magenta) and γ-tubulin (green) is shown in the CP epithelial cells in newborn *Gemc1*^*−/−*^ and wild type animals. Boxed regions are shown in higher magnification on the right. DAPI staining (cyan) labels nuclei. Scale bar, 20 μm. BB, basal body. (*C*, *D*, *E*) Immunohistochemical analyses of the expression of OTX2 (*C*), AQP1 (*D*), and TTR (*E*) are shown at days E13.5 (C) and P7 (*C*, *D*, *E*) in the roof plate (*C*, marked by dotted lines) and CP in *Gemc1*^*−/−*^ (arrowheads) and wild type (arrows) animals. Scale bars, 50 μm.

**Figure 2.**
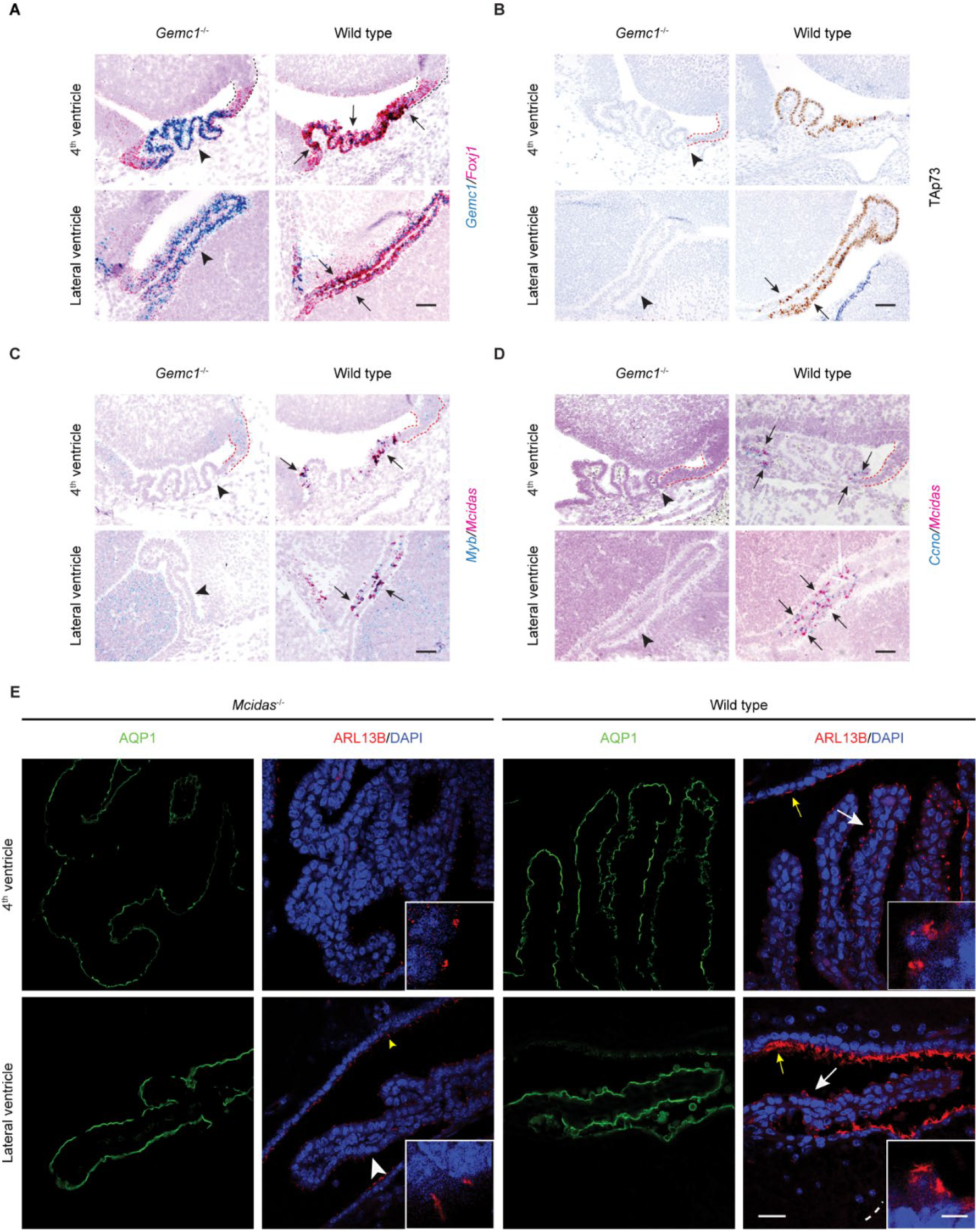
Defective multiciliation network in *Gemc1*-null CP. (*A*) RNAscope analysis of the expression of *Gemc1* and *Foxj1* is shown at day E13.5 in roof plate (marked by dotted lines) and CP in *Gemc1*^*−/−*^ (arrowheads) and wild type (arrows) animals. Scale bar, 50 μm. (*B*) The expression of TAp73 is shown at day E13.5 in roof plate (marked by red dotted lines) and CP in *Gemc1*^*−/−*^ (arrowheads) and wild type (arrows) animals. Scale bar, 50 μm. (*C*, *D*) RNAscope analysis of the expression of *Mcidas* and *Myb* (*C*), or *Mcidas* and *Ccno* (*D*) is shown at day E13.5 in roof plate (marked by dotted lines) and CP in *Gemc1*^*−/−*^ (arrowheads) and wild type (arrows) animals. Scale bar, 50 μm. (*E*) The expression of ARL13B (red) and AQP1 (green) is shown in CP epithelial cells. Cilia in newborn *Mcidas*^*−/−*^ (white arrowheads) and wild type (white arrows) animals are magnified in inset pictures. Ependymal cells lining the ventricles are shown in *Gemc1*^*−/−*^ (yellow arrowheads) and wild type (yellow arrows) animals. DAPI staining (blue) labels nuclei. Scale bars, 25 μm, 5 μm (inset).

Importantly, the expression of *Mcidas*, *Myb* and *Ccno* is upregulated in a subpopulation of CP epithelial cells next to the roof plate in wild type animals at day E13.5, whereas their expression is lost in the absence of *Gemc1* (Fig. 2C,D), suggesting that GEMC1 activates a MCIDAS-dependent program that mediates multiciliogenesis in the CP during embryonic development. Indeed, solitary primary cilia were detected in *Mcidas*^−/−^ CP epithelial cells (Fig. 2E). *Mcidas*-deficient CP exhibited OTX2 and AQP1 expression similar to that of wild type CP (Fig. 2E; Supplemental Fig. S4A,B). Further, the expression of *Gemc1*, its transcriptional targets *Foxj1* and TAp73 remained unaltered by *Mcidas* loss, though *Mcidas* overexpression was able to stimulate TAp73 expression (Supplemental Fig. S3C,D; Supplemental Fig. S4C,D). Taken together, these results indicate that MCIDAS functions in GEMC1-directed transcriptional network and plays an essential role in multiciliogenesis in the CP.

### SHH and NOTCH pathways drive CPC

We previously used a *Rosa26-NICD1* mouse strain that exhibits Cre-mediated expression of the intracellular domain of NOTCH1 (NICD1) and green fluorescent protein (GFP) (Murtaugh et al. 2003). After crossing with *Lmx1a-Cre* transgenic mice that express *Cre* in the roof plate/CP (Chizhikov et al. 2006), *Lmx1a-Cre;Rosa26-NICD1* (*Lcre;NICD1*) mice develop CPP that undergoes proliferation driven by SHH transiently expressed in the CP epithelium (Li et al. 2016). Loss of SHH signals in the CP led to cell cycle exit in tumor cells after birth, whereas treatment of tumor cells with recombinant SHH *in vitro* restored the proliferation of tumor cells. Further studies revealed that singular primary cilia of NOTCH-driven CP tumor cells critically mediates SHH signaling in tumor cells (Li et al. 2016).

Abnormal SHH and NOTCH pathway activities occur in human CP tumors (Li et al. 2016). To determine whether aberrant SHH signaling collaborates with NOTCH to drive aggressive CP tumors, we bred a mouse strain carrying a *Patched1* conditional allele (*Ptch*^*flox/flox*^) to *Lcre;NICD1* mice. Loss of *Ptch1* would constitutively activate SHH signaling in the roof plate/CP in *Lmx1a-Cre;Ptch*^*flox/flox*^;*NICD1* (*Lcre;Ptch*^*cko*^;*NICD1*) animals (Uhmann et al. 2007). Though most *Lcre;NICD1* mice survived normally, *Lcre;Ptch*^*cko*^;*NICD1* animals died perinatally from abnormal growth in the brain (Fig. 3A). At day E14.5, these animals displayed a dramatically enlarged roof plate, abnormal CP growth with increased cellularity, and loss of monolayer epithelial architecture (Fig. 3A; Supplemental Fig. S5A). The percentage of Ki-67^+^ cells in the CP of *Lcre;Ptch*^*cko*^;*NICD1* mice was significantly increased (Fig. 3B; *Lcre;NICD1* mice: 52.75 ± 2.72%, *n* = 4; *Lcre;Ptch*^*cko*^;*NICD1* mice: 64.86 ± 1.65%, *n* = 7; two-tailed unpaired *t*-test, *P* < 0.0001). Like NOTCH-driven CPP, abnormal CP growth in *Lcre;Ptch*^*cko*^;*NICD1* animals exhibited elevated expression of SHH pathway targets *Gli1* and *Mycn,* and reduced *Shh* expression (Supplemental Fig. S5B). Despite the expression of OTX2, CP tumors in *Lcre;Ptch*^*cko*^;*NICD1* and *Lcre;NICD1* animals showed reduced expression of AQP1, TTR and cytokeratins (Fig. 3B; Supplemental Fig. S5C). Notably, a small population of CP epithelial cells are mixed with tumor cells in these animals, possibly due to incomplete Cre-mediated activation of *NICD1* (Fig. 3B; Supplemental Fig. S5C). Together, these results indicate that aberrant SHH and NOTCH signaling can drive a malignant CP tumor that matches closely CPC in humans.

**Figure 3.**
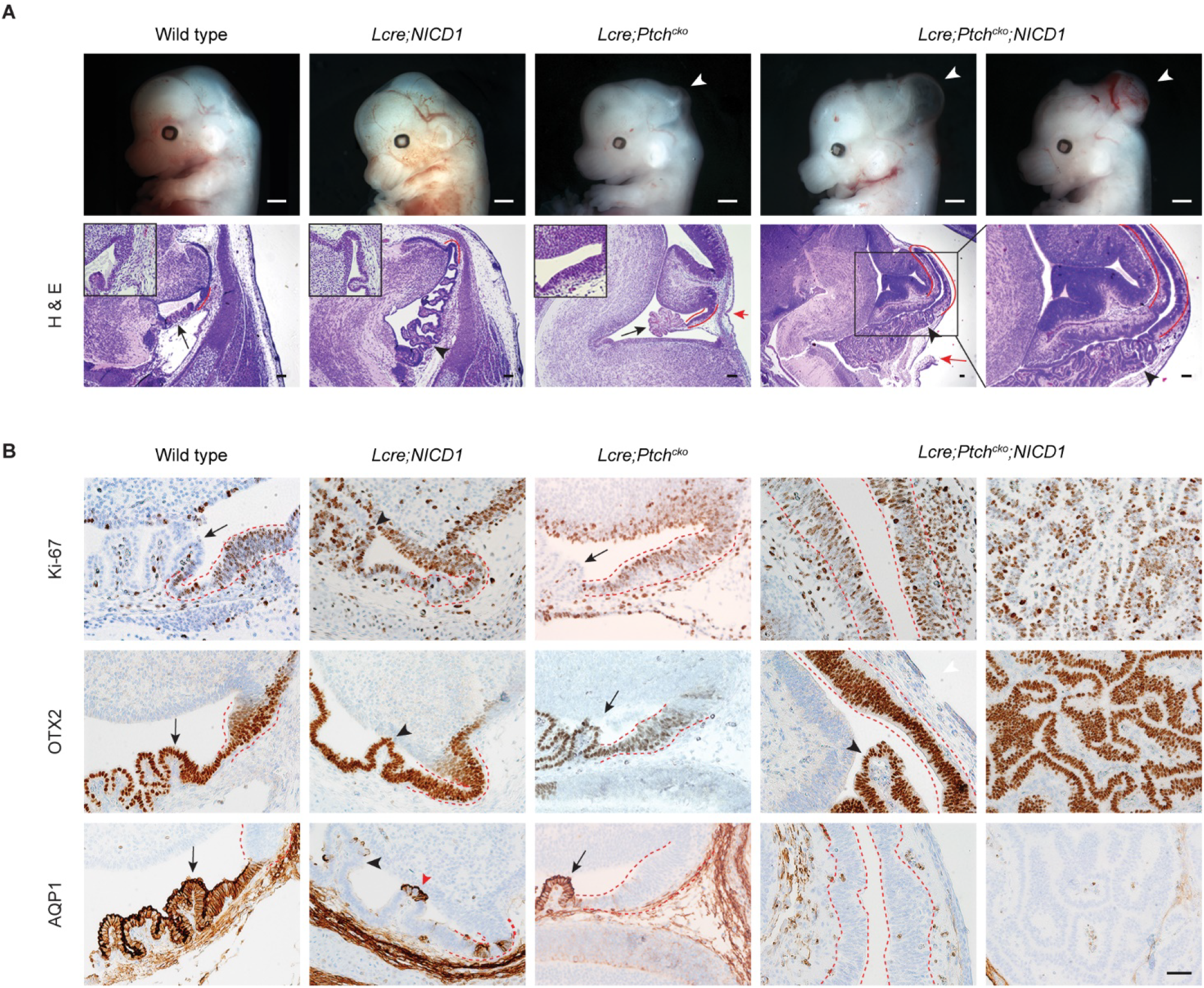
Aberrant NOTCH and SHH signaling drive CP carcinoma in mice. (*A*) Wild type, *Lcre;NICD1*, *Lcre;Ptch^cko^*, and *Lcre;Ptch*^*cko*^;*NICD1* animals are shown at day E14.5. Notice the cranium defects resulting from enlarged and folded roof plate in the midbrain-hindbrain region of *Lcre;Ptch*^*cko*^ and *Lcre;Ptch*^*cko*^;*NICD1* animals (white arrowheads). Hematoxylin and eosin (H&E) staining is shown of hindbrain roof plate (marked by red lines) and CP (black arrows) in wild type and *Lcre;Ptch*^*cko*^ animals, and abnormal CP growth (black arrowheads) in *Lcre;NICD1* and *Lcre;Ptch*^*cko*^;*NICD1* animals. Enlarged roof plate disrupts the cranium in *Lcre;Ptch*^*cko*^ and *Lcre;Ptch*^*cko*^;*NICD1* animals (red arrows). Red lines mark the roof plate magnified in inset images. Boxed region of hindbrain roof plate/CP in a *Lcre;Ptch*^*cko*^;*NICD1* animal is shown in higher magnification on the right. Scale bars, 100 μm. (*B*) The expression of Ki-67, OTX2, and AQP1 is shown in roof plate (marked by dotted lines) and CP (arrows) in wild type and *Lcre;Ptch*^*cko*^ animals, and tumor cells (black arrowheads) in *Lcre;NICD1* and *Lcre;Ptch*^*cko*^;*NICD1* animals at day E14.5. Residual AQP1-expressing epithelial cells (red arrowhead) are mixed with tumor cells in *Lcre;NICD1* animals. Scale bar, 50 μm.

### NOTCH activation leads to reduced multiciliation in CP tumors

In contrast to MCCs in the CP, NOTCH-driven CP tumor cells were monociliated and displayed increased expression of NOTCH targets *Hes1* and *Hes5* (Supplemental Fig. S5D; Supplemental Fig. S6A) (Li et al. 2016). Given the role of SHH signals in tumor cell proliferation, NOTCH might mediate the reduced multiciliation in CP tumors. To address this, tumor cells from *Lcre;NICD1* mice were cultured in the presence of a recombinant amino-terminal fragment of SHH (ShhN) and infected with viruses expressing GFP fused to dominant negative Mastermind-like protein 1 (dnMAML1) that disrupts MAML1 recruitment to the NOTCH transcriptional complex (Chang et al. 2011). Alternatively, tumor cells were treated with a small molecule Inhibitor of Mastermind Recruitment 1 (IMR-1) or its metabolite IMR-1A to disrupt NOTCH-mediated transcription (Astudillo et al. 2016). Remarkably, staining with the cilia markers ADP-ribosylation factor-like 13b (ARL13B) and γ-tubulin revealed multiple cilia in tumor cells within 72 hours after infection or IMR-1/IMR-1A treatment (Fig. 4A,B; Supplemental Fig. S6B; abundance of multiciliated GFP^+^ tumor cells: vehicle: 0 ± 0%, *n* = 4; IMR-1: 12.01 ± 1.34%, *n* = 3, *P* < 0.001; IMR-1A: 13.93 ± 1.97%, *n* = 3, *P* < 0.01, two-tailed unpaired *t*-test). IMR-1 treatment also markedly reduced tumor cell proliferation, whereas the expression of *Foxj1* was significantly increased by IMR-1 (Fig. 4C-E; Supplemental Fig. S6C). Importantly, after a 7-day IMR-1 treatment from day E10.5, multiciliated NICD1/GFP^+^ tumor cells were detected in *Lcre;NICD1* and *Lcre;Ptch*^*cko*^;*NICD1* animals, respectively (Fig. 4F; Supplemental Fig. S6D). IMR-1 treatment also significantly decreased tumor cell proliferation and tumor burden (Fig. 4G,H; Supplemental Fig. S6E). Together, these results demonstrate that aberrant NOTCH signaling impairs multiciliation that can be rescued by NOTCH inhibition, leading to cell cycle exit and reduced tumor growth.

**Figure 4.**
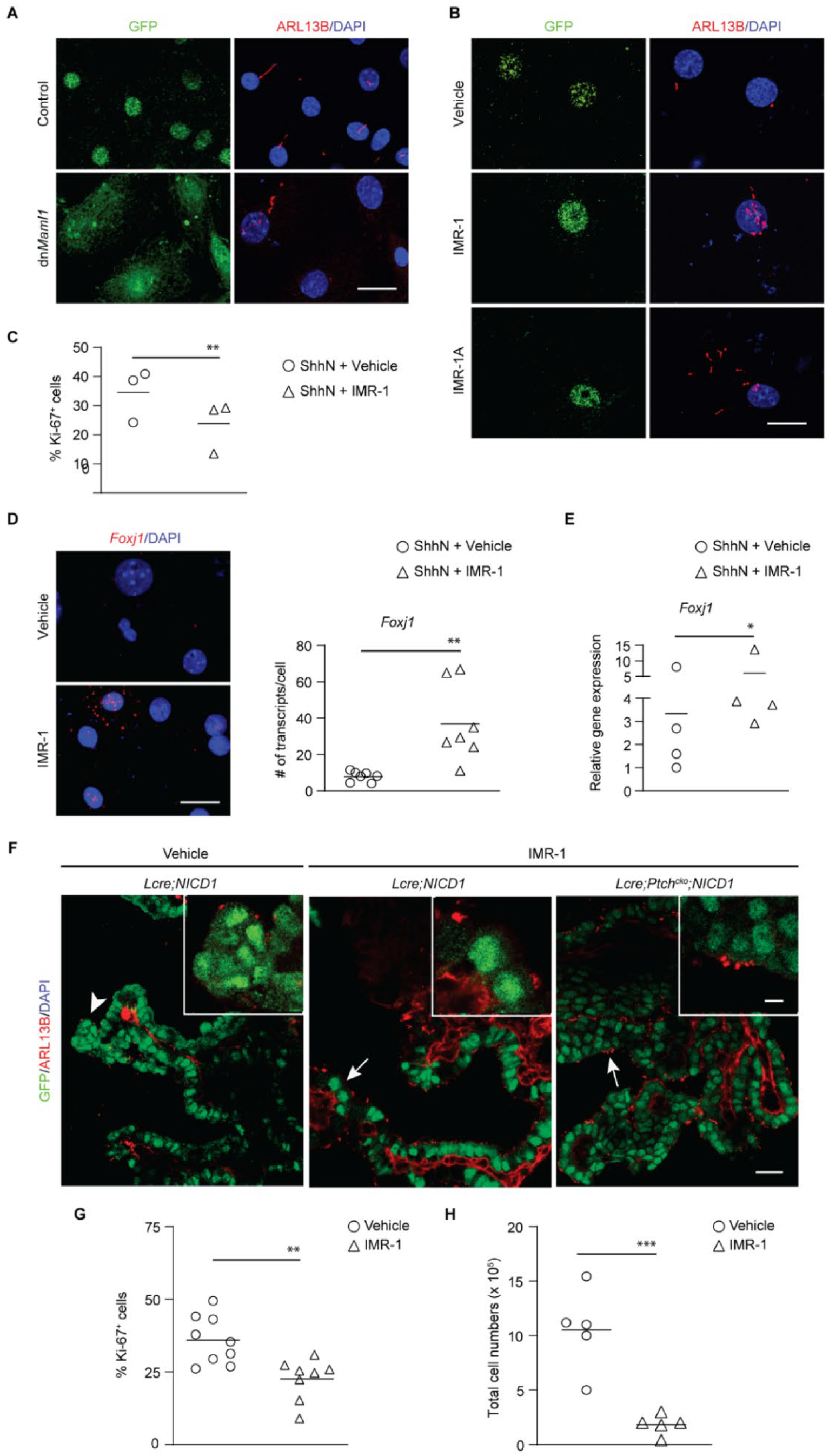
NOTCH activation leads to reduced multiciliation in CP tumors. (*A*, *B*) The expression of ARL13B (red) is shown in tumor cells infected with viruses expressing GFP-dnMAML1 or GFP (a), or treated with vehicle, or IMR-1/IMR-1A (b). GFP (green) labels infected or treated cells. DAPI staining (blue) labels nuclei. Scale bars, 20 μm. (*C*) The percentage of Ki-67^+^ cells is shown in NICD1^+^/GFP^+^ tumor cells treated with vehicle or IMR-1 (*n* = 3 animals per treatment; mean ± s.e.m., paired *t*-test, \*\**P* < 0.01). (*D*) RNAscope analysis of *Foxj1* expression (red) is shown in tumor cells treated with vehicle or IMR-1. DAPI staining (blue) labels nuclei. Scale bar, 20 μm. Quantification of mRNA transcripts is shown (*n* = 7 cells per treatment; mean ± s.e.m., two-tailed unpaired *t*-test, \*\**P* < 0.01). (*E*) RT-qPCR analysis of tumor cells treated with vehicle or IMR-1 (*n* = 4 animals per treatment; mean ± s.e.m., paired *t*-test, \**P* < 0.05). (*F*) The expression of ARL13B (red) is shown in NICD1^+^/GFP^+^ (green) tumor cells from *Lcre;NICD1* and *Lcre;Ptch*^*cko*^;*NICD1* animals treated with vehicle or IMR-1. Primary cilia in tumor cells treated with vehicle (arrowhead) or IMR-1 (arrows) are magnified in inset pictures. Scale bars, 20 μm, 5 μm (inset). (*G*, *H*) The percentage of Ki-67^+^ cells in NICD1^+^/GFP^+^ tumor cells (*G*) and total tumor cell numbers (*H*) are shown in *Lcre;NICD1* animals treated with vehicle or IMR-1 (g: *n* = 9 animals for vehicle, *n* = 8 animals for IMR-1; h: *n* = 5 animals per treatment; mean ± s.e.m., two-tailed unpaired *t*-test, \*\**P* < 0.01; \*\*\**P* < 0.001).

### Gemc1 suppression by NOTCH mediates defective multiciliation in CP tumors

Inhibition of the NOTCH pathway led to multiciliated differentiation in CP tumor cells; however, the underlying mechanisms of MCC differentiation regulated by NOTCH remains unclear. Reduced *Mcidas* and *Foxj1* levels from RNAseq studies of NOTCH-driven CPP (Fig. 5A) (Li et al. 2016), together with increased *Foxj1* levels following NOTCH blockade, suggests an interaction of NOTCH oncogenic signals with molecular apparatus of MCC differentiation (Kyrousi et al. 2015; Zhou et al. 2015; Lewis and Stracker 2020). Commensurate with this, NOTCH-driven CPP exhibited a transient increase in the levels of *Gmnn,* which is normally associated with proliferation and antagonizes GEMC1 functions in multiciliation (Fig. 5B) (Terre et al. 2016; Ortiz-Alvarez et al. 2019). In agreement with RNAseq, results from quantitative PCR, *in situ* hybridization and RNAscope studies all revealed decreased expression of *Gemc1*, *Mcidas,* their downstream targets *TAp73,* as well as *Foxj1* in NOTCH-driven CPPs and CPCs (Fig. 5A,B; Supplemental Fig. S7A-D). Moreover, immunofluorescence analysis showed that the expression of FOXJ1 and TAp73 was significantly reduced or lost in NOTCH driven CP tumor cells (Supplemental Fig. S7E,F). Together, these results demonstrated that the GEMC1-MCIDAS signaling is profoundly repressed in NOTCH-driven CP tumors.

**Figure 5.**
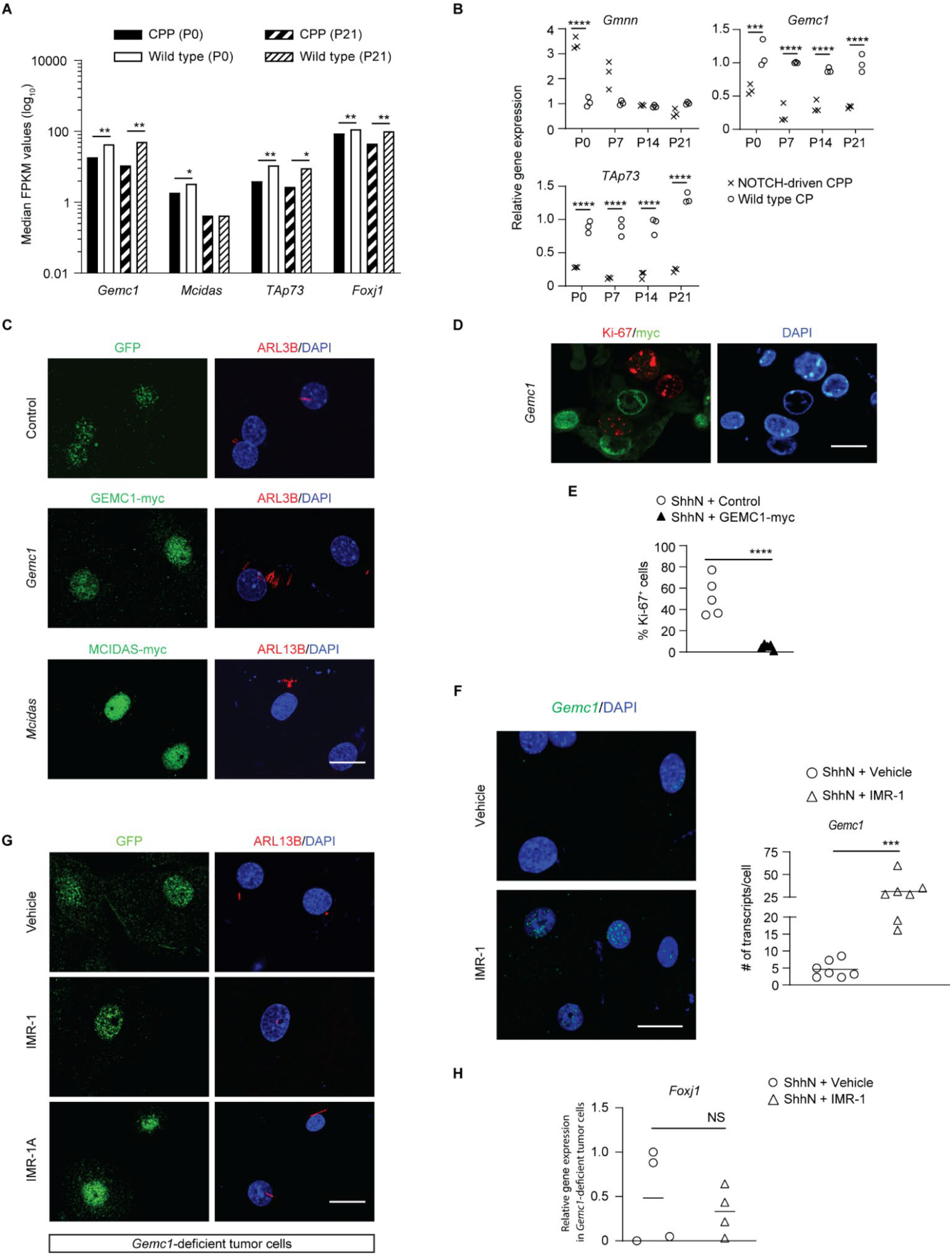
*Gemc1* suppression by NOTCH mediates multiciliation defects in CP tumors. (*A*) Median FKPM (fragments per kilobase of exon per million reads mapped) values of genes in NOTCH-driven CP tumors and wild type CPs (*n* = 3 specimens per time point, mean ± s.e.m., two-way ANOVA, \**P* < 0.05; \*\**P* < 0.01). (*B*) RT-qPCR analysis of NOTCH-driven CP tumors and wild type CPs (*n* = 3 animals per time point, mean ± s.e.m., two-tailed unpaired *t*-test, \*\*\**P* <0.001, \*\*\*\**P* < 0.0001). (*C*, *D*) The expression of ARL13B (*C*, red) and Ki-67 (*D*, red) is shown in tumor cells infected with viruses expressing GEMC1-myc, MCIDAS-myc, or GFP only. GEMC1-myc (green), MCIDAS-myc (green), or GFP (green) labels infected cells. DAPI staining (blue) labels nuclei. Scale bars, 20 μm. (*E*) The percentage of Ki-67^+^ cells in tumor cells infected with viruses expressing GEMC1-myc or GFP only (*n* = 5 animals per treatment, mean ± s.e.m., paired *t*-test, \*\*\*\**P* < 0.0001). (*F*) RNAscope analysis of *Gemc1* expression (green) is shown in tumor cells treated with vehicle or IMR-1. DAPI staining (blue) labels nuclei. Scale bar, 20 μm. Quantification of mRNA transcripts is shown (*n* = 7 cells per treatment; mean ± s.e.m., two-tailed unpaired *t*-test, \*\*\**P* < 0.001). (*G*) The expression of ARL13B (red) is shown in *Gemc1*-deficient tumor cells treated with vehicle or IMR-1/IMR-1A. GFP (green) labels tumor cells. DAPI staining (blue) labels nuclei. Scale bar, 20 μm. (*H*) RT-qPCR analysis of *Gemc1*-deficient tumor cells treated with vehicle or IMR-1 (*n* = 5 animals per treatment, mean ± s.e.m., two-tailed unpaired *t*-test, NS, not significant).

To understand the role of *Gemc1-Mcidas* suppression in defective multiciliation of CP tumors, myc-tagged GEMC1 or MCIDAS was expressed in tumor cells from *Lcre;NICD1* mice using viral vectors. Overexpression of either *Gemc1* or *Mcidas* led to the formation of multiple cilia and cell cycle withdrawal in tumor cells within 72 hours after infection (Fig. 5C-E; Supplemental Fig. S8A,B), thereby phenocopying NOTCH inhibition. Furthermore, IMR-1 treatment relieved suppression of *Gemc1* in tumor cells (Fig. 5F). To determine the role of *Gemc1* in multiciliation induced by NOTCH inhibition in CP tumor cells, we eliminated *Gemc1* by crossing a mouse strain hemizygous for a null and conditional allele (*Gemc1*^*−/flox*^) to *Lcre;NICD1* animals. Solitary cilia were retained in tumor cells from the resulting *Lcre;NICD1;Gemc1*^*−/flox*^ mice irrespective of IMR-1/IMR-1A treatment, whereas the proliferation of *Gemc1*-deficient tumor cells remained unaltered by NOTCH inhibition (Fig. 5G; Supplemental Fig. S8C,D; % of Ki-67^+^ cells in *Gemc1*-deficient tumor cells 72 hours after treatment: vehicle: 38.90 ± 1.44%, *n* = 6; IMR-1: 36.37 ± 4.65%, *n* = 6, two-tailed unpaired *t*-test, not significant). Accordingly, *Foxj1* mRNA levels in these tumor cells failed to be upregulated by IMR-1 (Fig. 5H), while reintroduction of *Gemc1* led to increased *Foxj1* expression in tumor cells from *Lcre;NICD1;Gemc1*^*−/flox*^ animals (Supplemental Fig. S8E). Therefore, these results indicate that monociliation in CP tumors is maintained through NOTCH-mediated suppression of GEMC1-MCIDAS signaling cascade, whereas impairment of the GEMC1-dependent MCC program prevents the rescue of multiciliation defects by NOTCH inhibition.

### CPCs in humans exhibit reduced multiciliation and a deficient GEMC1 program

Most CP tumors in humans, especially CPCs, are comprised of monociliated tumor cells. Importantly, these CP tumors frequently display large-scale genomic alterations (Ruland et al. 2014; Japp et al. 2015; Merino et al. 2015; Li et al. 2016). Among them, recurrent chromosomal changes affect loci encompassing multiciliogenesis regulators, including *GEMC1* on chromosome 3 that is lost in all hypodiploid CPCs, *MCIDAS*, *CCNO*, microRNA 449 (*MIR449*) and *CDC20B* within the same locus of chromosome 5, and *MYB* on chromosome 6 that are lost in many CPCs (Fig. 6A). Conversely, *N*-acetyl galactosamine-type *O*-glycosylation enzyme *GALNT11*, a positive regulator of NOTCH signaling on chromosome 7, is gained in >80% CP tumors (Fig. 6A) (Boskovski et al. 2013; Ruland et al. 2014; Japp et al. 2015; Merino et al. 2015; Lewis and Stracker 2020). In agreement, most human CPCs display significantly reduced or complete loss of *GEMC1* expression, and to a lesser degree, decreased *FOXJ1* expression (Fig. 6B; Table 1). In these tumors, *GEMC1* expression is heterogeneous and only detected in a subpopulation of tumor cells. *GEMC1* and *FOXJ1* expression in CPP follows a similar trend (Fig. 6B; Table 1). Interestingly, TAp73 expression in human CP tumors varies from significantly reduced to normal levels (Supplemental Fig. S9A). Moreover, analysis of a published dataset revealed differential expression of genes involved in ciliogenesis in human CP tumors, contributing to significant enrichment of the pathway (Supplemental Fig. S9B,C) (Merino et al. 2015). Thus, CPCs in humans are characterized by cilia defects and a deficiency in the *GEMC1* transcriptional program.

**Table 1.** Summary of RNAscope study of gene expression in human CP tumors.

| Tumor type |  | < 10% tumor cells | 10 - 50% tumor cells | > 50% of tumor cells |
| --- | --- | --- | --- | --- |
| CPC | *GEMC1* | 72.7% (8/11) | 27.3% (3/11) | 0% (0/11) |
|  | *FOXJ1* | 63.6% (7/11) | 27.3% (3/11) | 9.1% (1/11) |
| CPP | *GEMC1* | 80.6% (25/31) | 16.1% (5/31) | 3.3% (1/31) |
|  | *FOXJ1* | 54.8% (17/31) | 32.3% (10/31) | 12.9% (4/31) |

CPC: 11 choroid plexus carcinomas from 11 individuals; CPP: 31 choroid plexus papillomas from 31 individuals.

**Figure 6.**
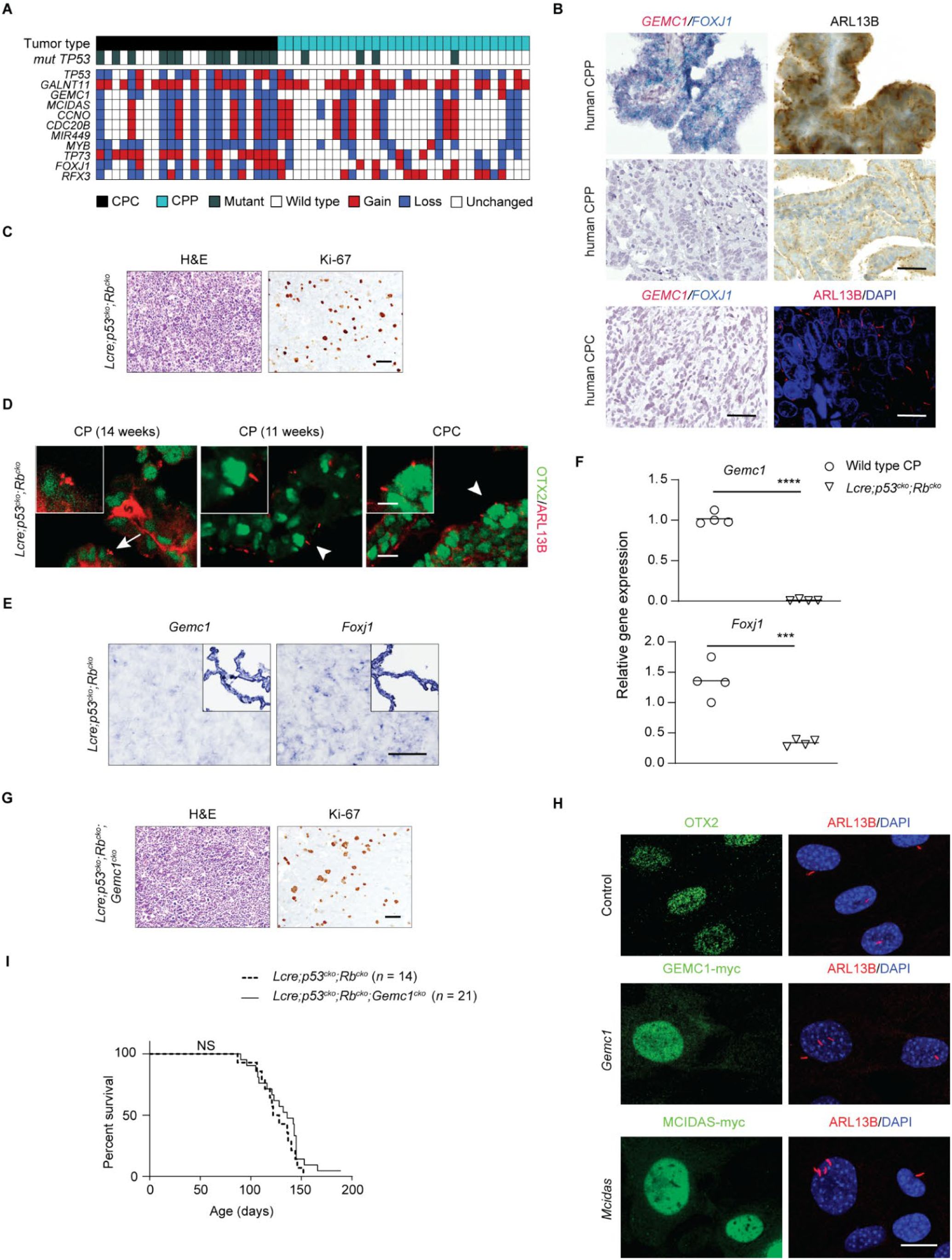
Multiciliation defects and GEMC1 program deficiencies in CP tumors. (*A*) Copy number analysis of human CPC (*n* = 23 tumors from 23 individuals) and CPP (*n* = 32 tumors from 32 individuals). (*B*) Representative images of human CP tumors showing *GEMC1* and *FOXJ1* expression by RNAscope (left panels), and ARL13B expression by immunostaining (brown signals in upper right panels, red signals in lower right panel). DAPI staining (blue, lower left panel) labels nuclei. Scale bars, 25 μm. (*C*) H&E staining and Ki-67 expression are shown in CPC from *Lcre;p53*^*cko*^;*Rb*^*cko*^ animals. Scale bar, 50 μm. (*D*) The expression of ARL13B (red) and OTX2 (green) is shown in the CP or CPC in *Lcre;p53*^*cko*^;*Rb*^*cko*^ animals. Cilia of multiciliated epithelial cells (arrow) or primary cilia of monociliated tumor cells (arrowheads) are magnified in inset pictures. DAPI staining (blue) labels nuclei. Scale bars, 20 μm, 5 μm (inset). (*E*) *In situ* hybridization of *Gemc1* and *Foxj1* mRNA in CPC from *Lcre;p53*^*cko*^;*Rb*^*cko*^ animals. Unaffected part of CP is shown in inset images. Scale bar, 25 μm. (*F*) RT-qPCR analysis of *Gemc1* and *Foxj1* expression in CPC and wild type CPs (*n* = 4 animals per tissue type, mean ± s.e.m., two-tailed unpaired *t*-test, \*\*\**P* <0.001, \*\*\*\**P* < 0.0001). (*G*) H&E staining and Ki-67 expression are shown in CPC from *Lcre;p53*^*cko*^;*Rb*^*cko*^;*Gemc1*^*cko*^ animals. Scale bar, 50 μm. (*H*) The expression of ARL13B (red) is shown in tumor cells from *Lcre;p53^cko^;Rb^cko^*;*Gemc1*^*cko*^ infected with viruses expressing GEMC1-myc or MCIDAS-myc. GEMC1-myc, or MCIDAS-myc (green) or OTX2 (green) labels tumor cells. DAPI staining (blue) labels nuclei. Scale bar, 20 μm. (*I*) Kaplan-Meier curve depicting the survival of *Lcre;p53*^*cko*^;*Rb*^*cko*^ and *Lcre;p53^cko^;Rb^cko^*;*Gemc1*^*cko*^ mice, NS, not significant.

### Gemc1 loss mediates cilia defects in CPCs with deficient Trp53 signaling

CPC frequently occurs in Li-Fraumeni syndrome patients, whereas homozygous somatic *TP53* mutations in sporadic CPC predict poor outcome (Tabori et al. 2010; Merino et al. 2015). In agreement, *Trp53* deletion combined with *Rb1* loss or *Myc* overexpression leads to murine CPC with characteristics of their human counterpart (Tong et al. 2015; El Nagar et al. 2018; Shannon et al. 2018; Merve et al. 2019; Wang et al. 2019). We crossed *Lmx1a-Cre* mice with a mouse strain carrying conditional alleles of *Trp53* and *Rb1* (*p53^flox/flox^;Rb^flox/flox^*) (Marino et al. 2000). All *Lmx1a-Cre;p53*^*flox/flox*^;*Rb*^*flox/flox*^ (*Lcre;p53*^*cko*^;*Rb*^*cko*^) mice developed CPC (Fig. 6C). At 11 weeks of age, OTX2^+^ monociliated cells were present among MCCs in the CP of *Lcre;p53*^*cko*^;*Rb*^*cko*^ animals (Fig. 6D). Like its human counterpart, CPC from these mice consisted of monociliated tumor cells, and exhibited markedly reduced *Gemc1* and *Foxj1* expression (Fig. 6D-F). Thus, *Trp53*-deficient CPC recapitulates the multiciliation defect and GEMC1 MCC program deficiency in human CPC.

To determine the role of *Gemc1* in multiciliation defects in CPC with deficient *Trp53* signaling, we subsequently targeted *Gemc1* by breeding *Lcre;p53*^*cko*^;*Rb*^*cko*^ and *Gemc1*^*−/flox*^ animals. Like *Lcre;p53*^*cko*^;*Rb*^*cko*^ mice, *Lcre;p53^cko^;Rb^cko^*;*Gemc1*^*−/flox*^ (*Lcre;p53*^*cko*^;*Rb*^*cko*^;*Gemc1*^*cko*^) mice developed CPCs that expressed OTX2 and were proliferative, whereas the expression of AQP1 was significantly reduced in these tumors (Fig. 6G; Supplemental Fig. S10A,B). In comparison with dramatic reduction in the expression of FOXJ1 in *Trp53*-deficient CPC, TAp73 expression was more variable (Supplemental Fig. S10A,B). In line with this, analysis of published microarray data from *Trp53*-deficient CPC showed reduced expression of genes present in adult CP, along with those involved in multiciliogenesis (Supplemental Fig. S10C,D) (Tong et al. 2015). Subsequently, Myc-tagged GEMC1 or MCIDAS was expressed in tumor cells from *Lcre;p53*^*cko*^;*Rb*^*cko*^;*Gemc1*^*cko*^ mice. Reintroduction of *Gemc1* or *Mcidas* into *Trp53*-deficient CPC activated the expression of *TAp73* and *Foxj1*, and rescued the cilia defect in these tumor cells by converting them into multiciliated post-mitotic cells (Fig. 6H; Supplemental Fig. S10E-G). Nonetheless, *Gemc1* loss failed to accelerate tumor development in *Lcre;p53*^*cko*^;*Rb*^*cko*^;*Gemc1*^*cko*^ animals (Fig. 6I). In addition, neither *Lcre;p53*^*flox/+*^;*Rb*^*cko*^;*Gemc1*^*cko*^ nor *Lcre;p53*^*cko*^;*Rb*^*flox/+*^;*Gemc1*^*cko*^ animals developed CPC (data not shown). In agreement, despite the suppression of *Gemc1-Micdas* signaling cascade by NOTCH oncogene, no CP tumor was detected after combined loss of *Gemc1* and *Patched1* in *Lcre;Ptch*^*cko*^;*Gemc1*^*−/flox*^ animals (data not shown), suggesting that loss of *Gemc1* transcriptional program and multicilia clusters in the CP are insufficient to drive tumorigenesis. Together, these data provide incriminating evidence that defective a GEMC1-MCIDAS transcriptional program mediates the cilia defect in *Trp53*-deficient CPC and promotes tumor growth.

## Discussion

The CP in each brain ventricle consists of a fibro-vascular core, encapsulated by epithelial MCCs (Narita and Takeda 2015; Fame and Lehtinen 2020). The CP plays an important role in safeguarding the environment in the central nervous system through its barrier functions and CSF secretion. Though the unique arrangement of multiple cilia serves no significant purpose in fluid flow, they have been suggested to contribute to chemosensory functions (Narita et al. 2010; Narita and Takeda 2015). Our data have revealed that GEMC1-MCIDAS activity is responsible for the differentiation of multiple cilia clusters of CP epithelial cells. Targeted disruption of either *Gemc1* or *Mcidas* led to a lack of cilia clusters in the CP and distinct alterations in the expression of downstream targets. As roof plate progenitors exit the cell cycle and commit to multiciliogenesis, TAp73 is turned on in multiciliated epithelial cells (Wildung et al. 2019). Interestingly, both *GEMC1* and *MCIDAS* were able to stimulate *TAp73* expression (Lalioti et al. 2019a). *Gemc1* is expressed in differentiated epithelial cells, where its loss prevented activation of downstream targets including *Mcidas*, *TAp73* and *Foxj1* (Lalioti et al. 2019a; Lewis and Stracker 2020). In contrast, the expression of *Mcidas*, *Myb* and *Ccno* is detected only in early multiciliated epithelial cells. Despite lack of MCCs in *Mcidas*-deficient CP, the expression of *Gemc1*, *TAp73* and *Foxj1* in the CP remained unaffected by *Mcidas* loss, consistent with the role of *Mcidas* in centriole amplification in MCC progenitors during the process of multiciliation (Tan et al. 2013; Boon et al. 2014; Ma et al. 2014; Wallmeier et al. 2014). The overlapping yet unique downstream targets of *Gemc1* and *Mcidas* in the CP are consistent with their distinct step-wise requirement in MCC differentiation in diverse epithelia (Lu et al. 2019).

Outcomes of CPC remain dismal, leaving patients vulnerable to devastating consequences (Gozali et al. 2012; Zaky and Finlay 2018). The development of safer and more effective therapies for CPC requires a better understanding of its biology. Though advances in genomic sequencing have driven the growth of precision cancer therapies, genetic drivers remain largely unknown in most CPCs. The frequency and scale of chromosomal alterations in these malignancies present a great challenge in the identification of driving events and actionable targets (Tong et al. 2015; Wang et al. 2019). The small patient population for this rare cancer and lack of experimental models further complicate drug screening efforts (El Nagar et al. 2018; Shannon et al. 2018; Merve et al. 2019). These daunting challenges demand alternative approaches to identify therapeutic strategies. Our data have provided evidence that multiciliation represents a barrier to tumorigenesis. Cross species molecular analyses have revealed consistent disruption of the multiciliogenesis network in CPC. The prevalence of solitary cilia suggested that CPC behaves like a ciliopathy, with the loss of or inability to generate cilia clusters in epithelial cells. Thus, therapeutic strategies aimed at restoring multiple cilia may suppress CP tumors.

Findings from this study have shed light on the dynamic interaction of the multiciliogenesis program with developmental signaling during CP differentiation and tumorigenesis. NOTCH signaling suppresses multiciliated differentiation of roof plate progenitors, thereby preserving primary cilium-based SHH signaling (Li et al. 2016). The enlarged roof plate in *Lcre;Ptch*^*cko*^;*NICD1* animals is consistent with the developmental origin and cilia defect of CPC driven by NOTCH and SHH signaling. These animals serve as ideal therapeutic models for congenital or infantile CP tumors, a rare condition associated with high morbidity and mortality (Crawford and Isaacs 2019; Toescu et al. 2019). Indeed, NOTCH inhibition by IMR-1 rescued the cilia deficit by forming multiple cilia and decreasing mitosis in tumor cells. These results illustrate the potential benefit of restoring multiciliation in CP tumor treatment. Several drugs targeting the NOTCH complex may provide additional options for this therapeutic approach (Hurtado et al. 2019).

The formation of MCC is inhibited by NOTCH signaling, the loss of which leads to supernumerary MCCs (Deblandre et al. 1999; Liu et al. 2007; Guseh et al. 2009; Tsao et al. 2009; Morimoto et al. 2010). The differentiation of MCCs requires inhibition of NOTCH pathway; however, it is unclear how NOTCH impacts MCC fate choice. Though previous studies showed that NOTCH decreases GEMC1-MCIDAS activity (Stubbs et al. 2012; Kyrousi et al. 2015), our results showed for the first time that NOTCH suppressed the expression of *Gemc1* and *Mcidas* to impair multiciliation in CP tumor cells. Furthermore, *Gemc1* expression was required for MCC differentiation following treatment by a NOTCH inhibitor, indicating that GEMC1-MCIDAS multiciliogenesis network is required downstream of NOTCH regulation and comprise a potent anti-tumor mechanism. *Mcidas* is capable of promoting multiciliogenesis during development independent of *Gemc1* (Lu et al. 2019). We also found that *Mcidas* could elicit the same multiciliation response as *Gemc1* in tumor cells. Although *Foxj1* is sensitive to *Gemc1* status in CP tumors, loss of TAp73 does not affect *Foxj1* expression in the CP as opposed to in the airway, indicating that *TAp73* is not entirely integrated in the *Gemc1*-*Foxj1* axis in the CP (Wildung et al. 2019). Consistently, variable TAp73 expression was present in *Trp53*-deficient CP tumors in mice regardless of *Gemc1* status, as well as CP tumors in humans. Further elucidation of the *Gemc1*-*Mcidas* regulatory cascade in the CP may help to identify critical regulators of multiciliogenesis that can be targeted for therapeutic development.

A deficient GEMC1 transcriptional program was also implicated in defective multiciliation in CPC driven by mechanisms independent of NOTCH. Gross chromosomal changes are a hallmark of CP tumors, whereas *TP53* mutations are present in a significant fraction of CPC in humans (Ruland et al. 2014; Japp et al. 2015; Merino et al. 2015). Frequent loss of chromosomes 3 and 5, along with reduced *GEMC1* expression, suggests a role for GEMC1-MCIDAS signaling in ciliation defect in this subset of CPC. NOTCH inhibition is predicted to be ineffective in these tumors. Yet, reintroduction of either *Gemc1 or Mcidas* promoted multiciliation and blocked tumor cell proliferation, indicating that loss of GEMC1-MCIDAS activity is essential for preventing multiciliated differentiation in *Trp53*-deficient CPC. In agreement, decreased FOXJ1 expression was observed in human CPCs (Abedalthagafi et al. 2016).

Given the reduced *GEMC1* expression we observed in human CPPs, and reduced multiciliation in *Myc*-driven CP tumors (Shannon et al. 2018; Wang et al. 2019), disruption of the *GEMC1-MCIDAS* regulation may account for multiciliogenesis defects in diverse CP tumor entities. Cilia defects represent early events to facilitate cell cycle progression in nascent tumor cells; however, forced monociliation through *Gemc1* loss is insufficient to replace NOTCH or *Trp53/Rb1* loss in murine knockout models. It is likely that *Gemc1* is among crucial targets of oncogenic signaling through transcriptional and post-transcriptional mechanisms. The challenges associated with the disruption of a highly complex multiciliogenesis process unique to specialized MCCs may explain CPC as a rare cancer that occurs most commonly in young children. Cross species analysis of syntenic regions of genomic changes may identify shared mechanisms (Tong et al. 2015). A recent study of CP tumors in dogs identified several regions syntenic to loci within chromosome losses in human CP tumors, including loci in the vicinity of *GEMC1* on chromosome 3q28, and *MCIDAS* on chromosome 5q11 (Ancona et al. 2018). Analysis of syntenic regions in CPC across species may identify their critical interactors and provide crucial insights into *GEMC1-MCIDAS* regulation and cilia defects in CPC.

GEMC1-MCIDAS network is essential for multiciliated differentiation of ependymal cells from radial glial cells, and antagonism between GMNN and GEMC1 balances ependymal and stem cell populations in adult neurogenic niche (Jacquet et al. 2009; Kyrousi et al. 2016; Terre et al. 2016; Lalioti et al. 2019b; Ortiz-Alvarez et al. 2019). Interestingly, FOXJ1-dependent program is enriched in PF-EPN-B subset of ependymoma, suggesting a transcriptional profile similar to the ependymal cells, whereas other ependymoma subgroups are more aligned with radial glial cells (Johnson et al. 2010; Witt et al. 2011; Abedalthagafi et al. 2016; Mack et al. 2018). Consistent with this, reduced or abnormal ciliogenesis occurs in ependymoma (Alfaro-Cervello et al. 2015), reminiscent of defective multiciliogenesis in CPC. Therefore, GEMC1-MCIDAS program may be suppressed to mediate multiciliation defects in ependymomas as well.

Overall, our study shows that the GEMC1-MCIDAS transcriptional program is required for MCC differentiation in the CP, whereas its compromise by oncogenic signals prevents multiciliation and promotes proliferation of CP tumor cells. Therefore, activation of multiciliogenesis may serve as a potential therapeutic strategy in a subset of CP tumors. However, the early events that lead to GEMC1-MCIDAS program activation remain poorly characterized, and detailed understanding of its regulation and molecular functions will be critical to facilitate the targeting of this pathway for the treatment of CPC.

## Materials and methods

### Animals

*Gt(ROSA)26Sor*^*tm1.Notch1Dam*^/*J* (*Rosa26-NICD1*) mice, B6N.129-*Ptch1*^*tm1Hahn*^/J (*Ptch*^*flox/flox*^) mice, B6.129P2-*Trp53*^*tm1Brn*^/J (*Trp53*^*flox/flox*^) mice, *Rb1^tm2Brn^*/J (*Rb*^*flox/flox*^) mice, and *C57BL/6* mice (all from Jackson Laboratory, Bar Harbor, ME), *Tg(Lmx1a-cre)1Kjmi* (*Lmx1a-Cre*) mice, *Gmnc*^*tm1Strc*^ (*Gemc1*^*−/+*^) mice and *Gmnc*^*tm1.1Strc*^ (*Gemc1*^*lox/+*^) mice were maintained by breeding with C57BL/6 mice. Animals were housed in the Animal Research Facility at New York Institute of Technology College of Osteopathic Medicine in accordance with NIH guidelines. All animal experimental procedures were approved by Institutional Animal Care and Use Committee (IACUC) and performed in compliance with national regulatory standards. *Mcidas* mutant mice were housed at the Biological Resource Center of the Agency for Science, Technology and Research (A*STAR) of Singapore and experiments performed with these animals followed guidelines stipulated by the Singapore National Advisory Committee on Laboratory Animal Research. All experimental procedures at the Institute for Research in Biomedicine were conducted following European and National Regulation for the Protection of Vertebrate Animals used for experimental and other scientific purposes (directive 86/609), internationally established 3R principles, and guidelines established by United Kingdom Coordinating Committee on Cancer Research. Experimental animals were administered 15 mg/kg IMR-1 (SML1812, Sigma-Aldrich, St. Louis, MO) or vehicle from day E10.5 through E16.5 (*Lcre;NICD1* mice: 13 animals for IMR-1, 14 animals for vehicle; *Lcre;Ptch*^*cko*^;*NICD1* animals: 5 animals for IMR-1, 4 animals for vehicle) by intraperitoneal injection of pregnant females.

### Human samples

CP specimens were procured with informed consent from patients following the requirements by institutional review boards at Shanghai East Hospital, Sanford Burnham Prebys Medical Discovery Institute, and University Medical Center Hamburg-Eppendorf. All CP specimens from Boston Children's Hospital were obtained under an approved institutional review board protocol. All tissues were handled in accordance with guidelines and regulations for the research use of human brain tissue set forth by the NIH (http://osp.od.nih.gov/o_ce-clinical-research-and-bioethics-policy). Diagnoses of human CP specimens from Boston Children's Hospital were reviewed by two neuropathologists (H.G.W.L., S. Santagata) using standard WHO criteria (Louis et al. 2007).

### Primary cell culture

Dissected CP specimens were dissociated using forceps under a stereoscope followed by enzymatic digestion at 37°C for 20 minutes with pronase (2 mg/ml, 537088, Calbiochem, San Diego, CA) in Hank’s balanced salt solution (HBSS, 14170-112; Life Technologies, Grand Island, NY) supplemented with 2 mM glucose. The Dulbecco’s modified Eagle’s medium/Nutrient Mixture F-12 Ham’s-Liquid Media (DMEM/F12, SH30261; HyClone Laboratories, Waltham, MA) supplemented with 10% fetal bovine serum (FBS, Atlanta Biologicals, Lawrenceville, GA) was added (1:1 ratio) to stop enzymatic digestion. Dissociated CP cells were centrifuged at 100 g for 2 minutes at 4°C. Cells were resuspended and cultured in DMEM/F12 supplemented with 10% FBS and 100U/mL penicillin/streptomycin (Life Technologies). Cytosine β-D-arabinofuranoside (Ara-C, 20 μM; C1768, Sigma-Aldrich) was added to culture media the next day to remove contaminating fibroblasts. Cultured CP cells were then treated with IMR-1 (25 μM) or IMR-1A (25 nM) (Sigma-Aldrich) in DMEM/F12 medium supplemented with ShhN (200 ng/ml) (Zhao et al. 2008), and B27 (Life Technologies). Dissociated tumor cells from *Lcre;NICD1* animals treated with IMR-1 or vehicle were resuspended in 0.2 ml cold phosphate-buffered saline (PBS) and spun onto a slide at 800 g for 5 min using a Shandon Cytospin 4 Cytocentrifuge (Thermo Scientific, Waltham, MA).

Primary CP tumor cells were not listed in the database of commonly misidentified cell lines maintained by ICLAC and NCBI Biosample. Analyses of gene expression, proliferation, and signal transduction were performed in primary CP tumor cells. Results from these studies and our previous report confirmed their identity. Given the short period of time (< 8 days) these cells in culture in the presence of antibiotics, tests were not performed for mycoplasma contamination.

### Viruses

cDNA encoding murine GEMC1 or MCIDAS with both Myc and FLAG tags at C-terminus (GEMC1-MycDDK, or MCIDAS-MycDDK) was amplified from plasmid pCMV6-Entry-*Gemc1*-MycDDK or pCMV6-Entry-*Mcidas*-MycDDK by PCR to introduce 5’ *Kpn I* and 3’ *Hind III* restriction sites. The PCR product was inserted into a pMiniT vector (E1202S, New England Biolab, Ipswich, MA). *KpnI - HindIII* fragment of *Gemc1-MycDDK* or *Mcidas-MycDDK* cDNA was then inserted into corresponding restriction sites in pShuttle-CMV vector (Agilent Technologies, Palo Alto, CA) to generate pShuttle-*Gemc1* or pShuttle-*Mcidas*, respectively. For dominant-negative form of MAML1 (*dnMaml1*), a pcDNA3-dnMaml1-GFP plasmid was used encoding the N-terminal NOTCH-binding domain of MAML1 fused in frame with GFP. The *dnMaml1-GFP* cDNA fragment was inserted into corresponding restriction sites in pShuttle-CMV to generate pShuttle-*dnMaml1*-GFP. All plasmids generated were verified by sequencing. The PmeI-linearized pShuttle-vectors carrying different cDNA fragments were introduced into the replication-deficient adenoviral vector pAdEasy-1 through homologous recombination in BJ5183 cells (Agilent Technologies). Successfully recombined adenoviral vector was verified by sequencing. Adenoviral plasmid was linearized by PacI digest and transfected into AD-293 cells (Agilent Technologies) to produce recombinant viral particles. Titers of purified recombinant viruses were determined by plaque assay and expressed as plaque-forming units (PFU) per milliliter (PFU/ml). Primary CP tumor cells were infected at 500 PFU/cell for 48 hours. All the procedures of production, purification and use of adenoviruses were approved by Institutional Biosafety Committee, performed in compliance with national standard for biosafety in microbiological and biomedical laboratories, and NIH Guidelines for Research involving Recombinant DNA Molecules.

### Histology

Animals were sacrificed and perfused with cold PBS followed by cold 4% paraformaldehyde (PFA). Tissue samples were dissected and fixed in 4% PFA overnight at 4°C. Fixed samples were processed for paraffin embedding and sectioning. Tissue sections were deparaffinized using CitriSolv (Decon Labs, King of Prussia, PA), and then rehydrated through graded ethanol solutions. For frozen tissues, fixed samples were further equilibrated in 20% sucrose in PBS for 24 - 48 hours at 4°C, then embedded in TissueTek-Optimal Cutting Temperature (O.C.T.) compound (Sakura Finetek, Torrance, CA) at −80°C. Frozen tissues were sectioned at 15 - 20 μm thickness on a cryostat. Cultured cells or acutely dissociated CP cells were fixed in 4% PFA for 10 minutes at room temperature.

### Immunohistochemistry, immunofluorescence, and immunocytochemistry

Immunostaining was carried out as previously described (Grausam et al. 2017). Tissue sections were counterstained with hematoxylin after immunohistochemistry. For immunofluorescence and immunocytochemistry, specimens were incubated with primary antibodies and then fluorescently labeled secondary antibodies (Jackson Immunoresearch, West Grove, PA). Stained specimens were cover-slipped with DAPI Fluoromount-G mounting medium (SouthernBiotech, Birmingham, Al). Primary antibodies used and dilution ratios are: mouse monoclonal anti-Acetylated α-Tubulin (1:500, ab24610, clone 6-11B-1, abcam, Cambridge, MA), mouse monoclonal anti-Acetylated α-Tubulin (1:500, T7451, clone 6-11B-1, Sigma-Aldrich), mouse monoclonal anti-ARL13B (1:500, clone N295B/66, NeuroMab, Davis, CA), rabbit anti-ARL13B (1:500, 17711-1-AP, Proteintech, Chicago, IL), mouse monoclonal anti-γ-Tubulin (1:10000, T6557, clone GTU-88, Sigma-Aldrich), chicken anti-GFP (1:1000, GFP-1010, Aves Lab, Tigard, OR), rabbit monoclonal anti-Ki-67 (1:100, clone SP6, ab16667, abcam), mouse monoclonal anti-Aquaporin 1 (1:1000, clone 1/22, ab9566, abcam), rabbit anti-Aquaporin 1 (1:1000, AB2219, EMD Millipore, Billerica, MA), and rabbit anti-OTX2 (1:500, AB9566, EMD Millipore), rabbit anti-Cytokeratins (1:100, Z0622, Dako, Carpinteria, CA), mouse monoclonal anti-FOXJ1 (1:50, 14-9965, Clone 2A5, eBioscience, San Diego, CA), and rabbit anti-TAp73 (1:200, ab40658, abcam), and sheep anti-Transthyretin (1:200, ab9015, abcam).

For primary cilia staining of CP specimens from mice, the cilia pattern was assessed by analyzing three distinct tissue regions of each sample. For staining of primary cilia in human CP tumors, tissues used include: normal CP: 1 disease-free individuals; CPP: 28 tumor specimens from 27 individuals; CPC: 16 tumors from 16 individuals. Cilia staining was assessed by analyzing 5 distinct tissue regions in each sample.

Human tissues used for TAp73 expression analysis include: normal CP: 4 individuals; CPP: 11 tumor specimens from 10 individuals; CPC: 10 tumors from 10 individuals. TAp73 expression was assessed in four distinct tissue regions: percent positive fields of view was calculated by scoring each 40 × field as “1” if there were Tap73^+^ cells present and “0” if there were none. The analysis was performed blinded by two independent investigators. For each tumor, the average of the scores for 5 fields of view is reported.

### Immunoblotting

Immunoblotting was carried out as described previously (Grausam et al. 2017). Briefly, tissue specimens were homogenized in lysis buffer and equal amounts (30 μg) of protein samples were separated by SDS-PAGE, and transferred to a polyvinylidene difluoride membrane. After incubation with 5% non-fat milk in PBST (PBS, 0.1% Tween 20) for 1 hour, the membrane was incubated with primary antibodies overnight at 4°C. After washing with PBST, the membrane was incubated for 1 hour with horseradish peroxidase (HRP)-conjugated secondary antibodies (GE Healthcare, Piscataway, NJ). The target protein on the membrane was detected by Pierce™ ECL Western Blotting Substrate (Thermo Scientific) or SuperSignal^®^ West Femto Maximum Sensitivity Substrate (Thermo Scientific) on Amersham Imager 680 (GE Healthcare). Primary antibodies used included: mouse monoclonal anti-β-Actin (1:1,000, clone AC-15, A5441, Sigma-Aldrich), mouse anti-FOXJ1 (1:1000, 14-9965, Clone 2A5, eBioscience), rabbit anti-Aquaporin 1 (1:1000, AB2219, EMD Millipore), rabbit anti-OTX2 (1:1000, AB9566, EMD Millipore), and rabbit anti-TAp73 (1:1000, ab40658, abcam), and goat anti-Myc (1:1000, ab9132, abcam).

### RNA isolation, RT-qPCR, in situ hybridization and RNAscope

Trizol and PureLink RNA Mini Kit (both from Life Technologies) were used to extract total RNA. For quantitative PCR following reverse transcription (RT-qPCR), total RNA samples were converted to cDNA using GoScript Reverse Transcription System (Promega, Madison, WI). All reaction conditions were set up in triplicate with ABsolute Blue QPCR Mix (Thermo Fisher Scientific) and then run on an ABI StepOne Plus Real-Time PCR System (Applied Biosystems, Foster City, CA). Gene-specific primers and probes were used (*Gmnn*: forward primer: 5’-TCTGCCAACAAGAGATCCATC-3’, reverse primer: 5’-GGAGCCCAAGAGAATGTGAAG-3’, probe: 5'-TGTCCCAAGGAGAACGCTGAAGATG-3'; *Gemc1*: forward primer 5’-ACCAGATTCTGACGTTGTAGTG-3’, reverse primer: 5’-GAGGCTATGGAAGATGCTGT-3’, probe: 5'-AGGAGGCCAGAGTTATAATTGCCCG-3'; *TAp73*: forward primer: 5’-AGCAATCTGACAGTACAACTTCT-3’, reverse primer: 5’-CATCCCTTCCAATACCGACTAC-3’, probe: 5'-TCACCTTCCAGCAGTCGAGCAC-3'). Data were analyzed using the ABI StepOnePlus Real‐Time PCR v2.0 software. Relative transcript levels were then calculated by the ΔΔCt method, and determined by the number of transcripts of genes of interest relative to those of an endogenous control β-actin (*Actb*, mouse) and normalized to the mean value of control samples. The value for each sample was obtained by averaging transcript levels of technical triplicates.

*In situ* hybridization was performed as described at *In Situ* Hybridization Core facility at Baylor College of Medicine (Yaylaoglu et al. 2005). Riboprobes for *Gli1*, *Mycn*, *Shh*, *Hes1*, *Hes5*, *Gemc1*, *Mcidas*, and *Foxj1* were used for CP specimens from wild type, *Lcre;Ptch^cko^*, *Lcre;NICD1*, and *Lcre;Ptch*^*cko*^;*NICD1* animals at day E14.5. Riboprobes for *Gemc1* and *Foxj1* were used for CP specimens from *Lcre;p53*^*cko*^;*Rb*^*cko*^ animals collected at terminal stage.

For RNAscope, *Gemc1* (510421), *Mcidas* (510401), *Myb* (510411), *Ccno* (546521) and *Foxj1* (317091) probes were used for CP specimens from wild type, *Gemc1*-deficient animals at days E13.5 and P7. *Gemc1*, *Mcidas*, and *Foxj1* probes were used for wild type CP, CP tumor from *Lcre;NICD1* mice, or NOTCH-driven CP tumor cells treated with IMR-1 in culture, or *Gemc1*-deficient NOTCH-driven CP tumor cells infected with viral vectors.

For human tissue samples, *GEMC1* (566231) and *FOXJ1* (430921) probes were used. RNA was visualized using RNAscope 2.5 HD Duplex Reagent Kit (322430, Advanced Cell Diagnostics, Hayward, CA) according to manufacturer's instructions. Human tissues used include: CPP: 31 tumor specimens from 31 individuals; CPC: 11 tumors from 11 individuals. *GEMC1* and *FOXJ1* expression was assessed in five distinct tissue regions: percentage of *GEMC1*-expressing cells or *FOXJ1*-expressing cells was calculated by averaging the numbers of *GEMC1*^+^ or *FOXJ1*^+^ cells per 100 cells from five distinct tissue regions of each specimen.

### Transmission electron microscopy

Tissues were fixed in 4% PFA, 1% glutaraldehyde in 0.1M cacodylate buffer (pH 7.4) at 4°C overnight. After washing with cacodylate buffer supplemented with 10% sucrose, tissues were post-fixed with 1% osmium tetroxide (OsO4), followed by incubation with 1% uranyl acetate in 30% ethanol. Tissue samples were dehydrated, then transferred to propylene oxide, and embedded in Eponate-12 epoxy resin (Ted Pella, Redding, CA). Tissue samples were sectioned (85nm thickness) with a Leica UC-6 ultramicrotome. Sections were counterstained with uranyl acetate and lead citrate, and observed under the JEOL 1400 transmission electron microscope (JEOL, Peabody, MA). Images were taken with a Gatan UltraScan 1000 CCD digital camera (Gatan, Pleasanton, CA).

### Image acquisition and quantitation

Whole-mount bright field was obtained using a Nikon SMZ1000 Stereomicroscope. Light and fluorescent microscopic images were obtained by a Nikon Eclipse 90i microscope system and a Nikon confocal microscope system A1^+^ (Nikon Instruments, Melville, NY). For analysis of cell proliferation in tissues, Ki-67^+^ cells in tumor cells or GFP^+^ cells were assessed from three distinct tissue regions for each animal. The percentage of Ki-67^+^ cells in the total tumor cell or GFP^+^ cell population was calculated by averaging the numbers of Ki-67^+^ cells per 100 tumor cells or GFP^+^ cells in all samples for each genotype at each time point or each treatment. For proliferation analysis of cultured cells, Ki-67 expression in tumor cells, GFP^+^ or GEMC1-myc^+^ cells was assessed by analyzing three distinct fields. The percentage of Ki-67^+^ cells was calculated by averaging the numbers of Ki-67^+^ cells per 100 tumor cells, GFP^+^ or GEMC1-myc^+^ cells of all samples for each treatment.

For analysis of multiciliation of cultured cells, primary cilia in tumor cells was assessed by analyzing ARL13B and γ-tubulin in three distinct fields. The percentage of multiciliated cells was calculated by averaging the numbers of multiciliated cells per 100 tumor cells of all samples for each treatment. The number of *Gemc1* or *Foxj1* mRNA transcripts detected by RNAscope was based on the fluorescent spot count of cells in 60 × microscopic images. The per-cell transcript copy number was determined by dividing the total spot count by the number of cells examined.

### Statistics

Multiple specimens were collected from independent samples or animals for each genotype. Both male and female animals were used at different time points. No randomization was used to determine how samples were allocated to experimental groups and processed. A group size of *n* = 10 (5 experimental, 5 control) will provide 90% power to detect a 22% change in assay results. Statistical analyses were performed with GraphPadPrism 8.0 (GraphPad Software Inc., La Jolla, CA). All pooled data were expressed as the mean ± standard error of the mean (SEM). Differences between two groups were compared using paired *t*-test or unpaired two-tailed *t*-test. Differences between multiple groups were analyzed with ANOVA followed by Tukey's multiple comparisons test. Results were considered significant at \**P* < 0.05; \*\**P* < 0.01; \*\*\**P* < 0.001; \*\*\*\**P* < 0.0001.

### Accession numbers

Published human CP tumor data sets (GSE14098, GSE60886), and data sets of murine *Trp53*-deficient CPC (GSE61659) were downloaded from GEO database for analysis. Hierarchical clustering was performed using Genesis (http://genome.tugraz.at/genesisclient/genesisclient_description.shtml). Pathway analysis using the GeneGoMetaCore Analytical Suite (http://genego.com; GeneGo) was used to score and rank pathways enriched in data sets by the proportion of pathway-associated genes with significant expression values. RNA-seq data (BioProject ID, PRJNA282889) were analyzed.

## Acknowledgements

We are grateful to Drs. Roger Packer, Huizhen Zhang, Brian Rood, William Weiss, Joanna Philips, Michael Taylor, James Loukides, and Sandro Santagata for providing human CP tumor samples, and Monica Calicchio for assisting with human specimens from Children’s Hospital (BCH). We thank Melanie Schweitzer, Amanda Chiang, Tamanna Sarowar, Brightlyn Kwa, Ibraheem Qureshi, Claire Evans, Phillip Petrasko, and Samuel Dooyema for technical assistance. The RNA In Situ Hybridization Core facility at Baylor College of Medicine is supported by an NIH Shared Instrumentation grant (1S10OD016167). This project was supported by NIH T32 HL110852 and BCH Faculty Development Fellowship (R.M.F.); NIH R01 NS088566 (MKL) and the New York Stem Cell Foundation (M.K.L.). M.K. Lehtinen is a New York Stem Cell Foundation – Robertson Investigator. U.S. is supported by the Fördergemeinschaft Kinderkrebszentrum Hamburg. T.H.S. was supported by the Spanish Ministry of Science, Innovation and Universities (MCIU: PGC2018-095616-B-I00/GINDATA and FEDER), the Centres of Excellence Severo Ochoa award and the CERCA Programme. S.R. is supported by funds from the A^*^STAR, Singapore. H.Z. is supported by New York Institute of Technology College of Osteopathic Medicine, Matthew Larson Foundation, NIH Institutional Development Awards (5P20GM103548 and 1P20GM103620-01A1) and NIH R01 CA220551.

## Author Contributions

Q.L., Z.H., Z. L., and H.Z. conceived and planned the project, and wrote the manuscript. H.G.W.L. reviewed diagnoses of human tissue samples. Q.L., N.S., Z.L., J.W., R.W.R. and U.S. analyzed morphology and gene expression of human specimens. R.M.F. and M.K.L. examined gene expression in human tissue samples. Q.L., Z.H., U.A., T.P., K.S., Z.L. and L.W. carried out morphological and gene expression studies in animal models. Y.H. produced and characterized viral vectors. P.C. conducted immunoblot analysis. Z.H. performed and analyzed results from cell culture experiments. T.Z. and A.A. performed histology and survival analysis. Z.H. conducted RNA-seq data analysis. H.L. and S.R. provided mutant mouse brains. B.T. and T.H.S. analyzed mutant mouse strain and T.H.S. and S.R. edited the manuscript.

