## Supplemental figures and legends for "GEMC1-MCIDAS transcriptional program regulates multiciliogenesis in the choroid plexus and acts as a barrier to tumorigenesis"

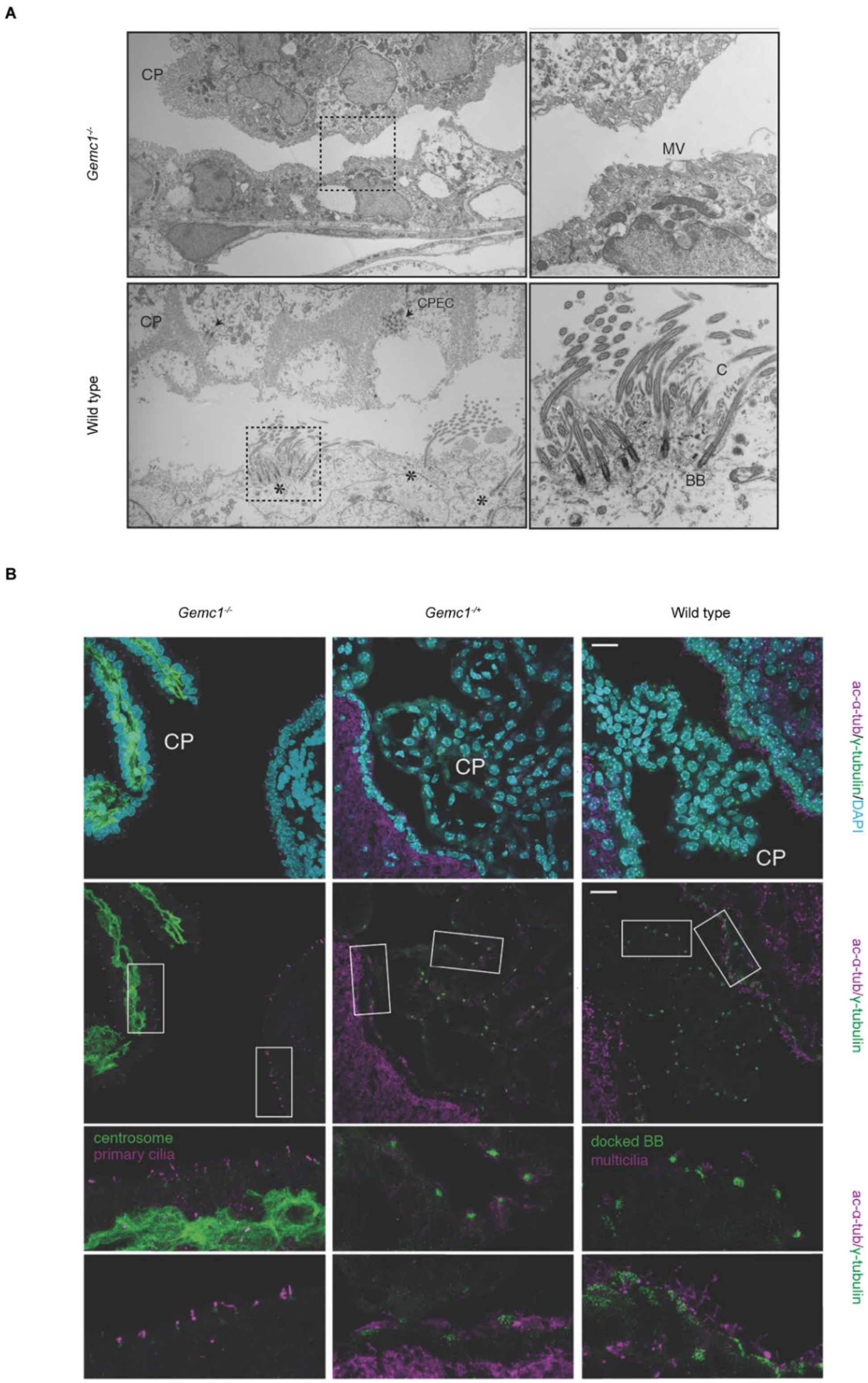


**Supplemental Figure S1. Loss of multiciliated cells in *Gemc1*-null brain.** (A) Transmission electron micrographs are shown of cilia clusters of CP epithelial cells (arrows) and motile cilia of ependymal cells (asterisks) at day P9 in *Gemc1*$^{-/-}$ and wild type animals. Boxed regions of the apical surface of ependymal cells are magnified on the right. CPEC, CP epithelial cells; BB, basal body; C, cilia; MV, microvilli. (B) The expression of acetylated α-tubulin (ac-α-tub, magenta) and γ-tubulin (green) is shown in the CP epithelium and ependyma at day P6 in *Gemc1*$^{-/-}$ and wild type animals. Boxed regions in the CP and ependyma are shown in higher magnification in lower two panels. DAPI staining (cyan) labels nuclei. BB, basal body. Scale bar, 20 µm.

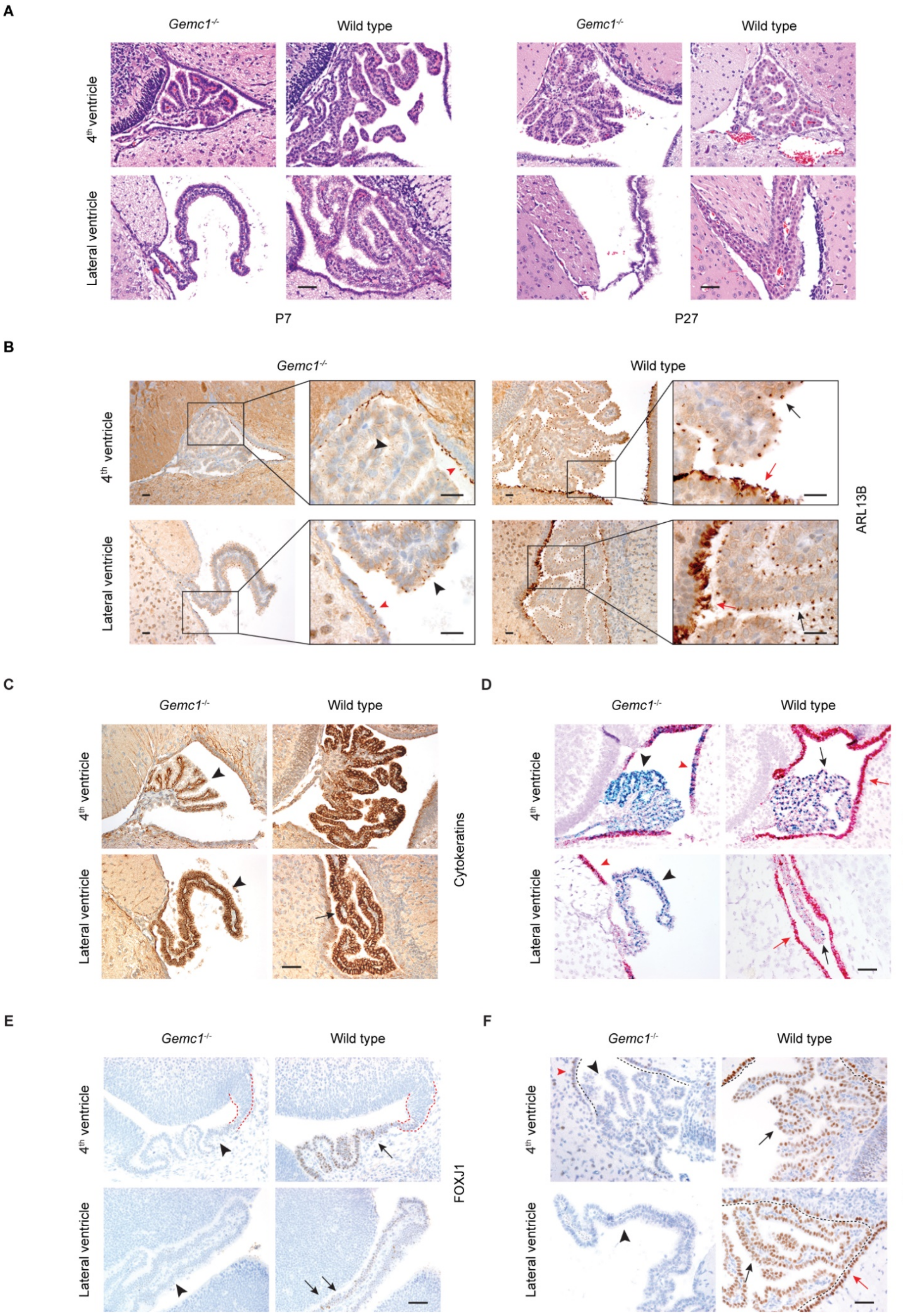


**Supplemental Figure S2**

**Supplemental Figure S2. Loss of MCCs and defective multiciliation network in *Gemc1*-null CP.** (A) H&E staining of CP in the 4$^{th}$ and lateral ventricles is shown at days P7 and P27 in *Gemc1*$^{-/-}$ and wild type animals. Scale bars, 50 µm. (B) Immunohistochemical analysis of ARL13B expression is shown at day P7. Boxed regions of the CP in *Gemc1*$^{-/-}$ (black arrowheads) and wild type (black arrows) animals are shown in higher magnification on the right. Ependymal cells lining the ventricles are shown in *Gemc1*$^{-/-}$ (red arrowheads) and wild type (red arrows) animals. Scale bars, 20 µm. (C) Immunohistochemical analysis of the expression of cytokeratins is shown at day P7 in the CP in *Gemc1*$^{-/-}$ (arrowheads) and wild type (arrow) animals. Scale bar, 50 µm. (D) RNAscope analysis of *Gemc1* and *Foxj1* expression is shown at day P7 in the CP in *Gemc1*$^{-/-}$ (black arrowheads) and wild type (black arrows) animals. Ependymal cells lining the ventricles are shown in *Gemc1*$^{-/-}$ (red arrowheads) and wild type (red arrows) animals. Scale bar, 50 µm. (E) The expression of FOXJ1 is shown at day E13.5 in roof plate (marked by dotted lines) and CP in *Gemc1*$^{-/-}$ (arrowheads) and wild type (arrows) animals. Scale bar, 50 µm. (F) The expression of TAp73 is shown at day P7 in the CP in *Gemc1*$^{-/-}$ (black arrowheads) and wild type (black arrows) animals. Ependymal cells lining the ventricles (marked by dotted lines) are shown in *Gemc1*$^{-/-}$ (red arrowhead) and wild type (red arrows) animals. Scale bar, 50 µm.

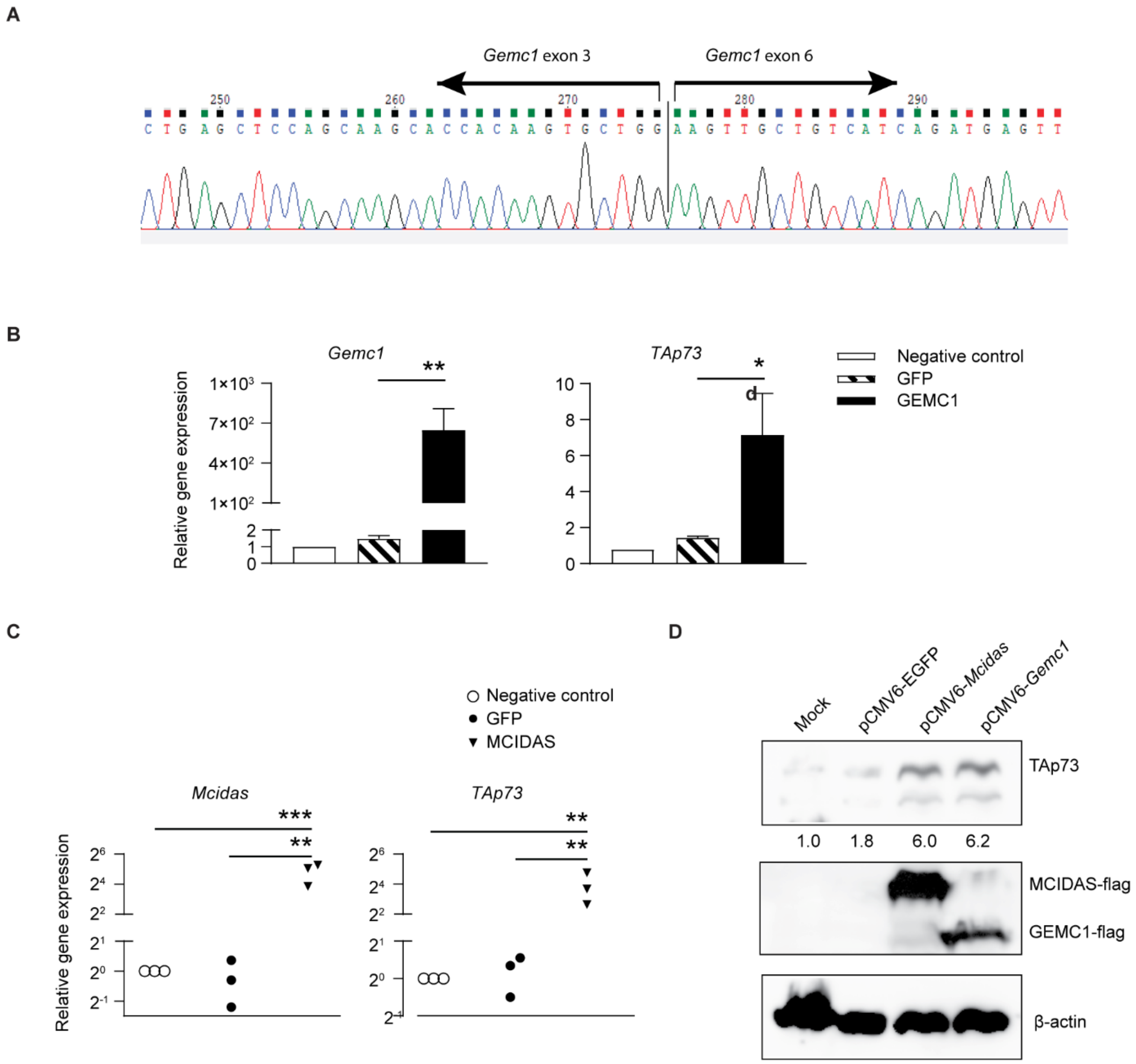


**Supplemental Figure S3. Analysis of *Gemc1*-driven gene expression.** (A) Sequencing trace image of *Gemc1* transcript in the CP of *Lcre;Gemc1*$^{-/flox}$ animals. Notice that exon 3 is spliced next to exon 6 in mutant transcript following the deletion of exons 4 and 5. (B) RT-qPCR analysis of mouse Inner Medullary Collecting Duct (mIMCD3) cells infected with viruses expressing GEMC1-myc or GFP only ($n$ = 3 samples per treatment, mean ± s.e.m., one-way ANOVA, *$P$ < 0.05; **$P$ < 0.01). (C) RT-qPCR analysis of mIMCD3 cells infected with viruses expressing MCIDAS-myc or GFP only ($n$ = 3 samples per treatment, mean ± s.e.m., one-way ANOVA, *$P$ < 0.05; **$P$ < 0.01). (D) Immunoblot analysis of HEK293T cells transfected with plasmids expressing FLAG-tagged MCIDAS or GEMC1, or GFP only. The value of each band indicates relative expression level normalized by loading control β-actin.

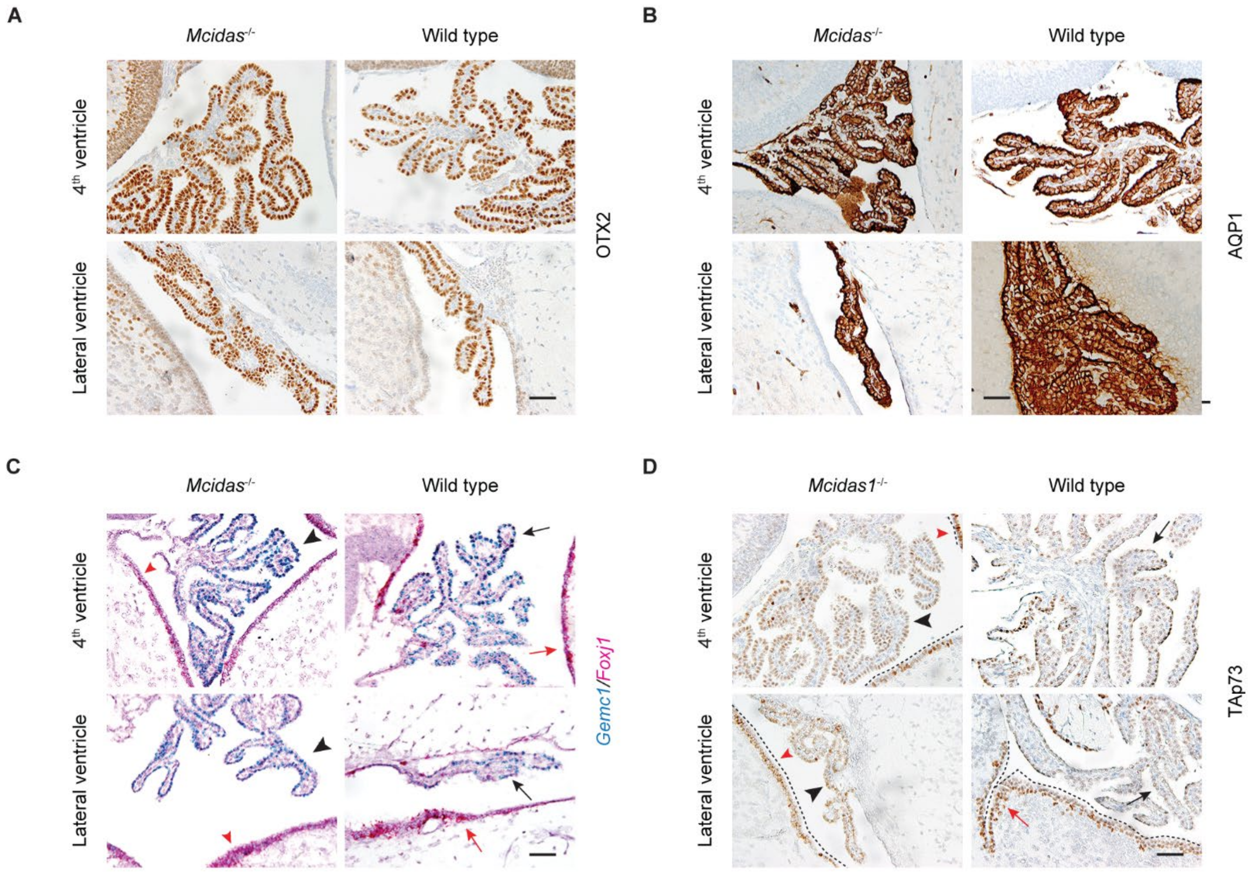


**Supplemental Figure S4. Defective multiciliation in *Mcidas*-null CP.** (A, B) Immunohistochemical analyses of the expression of OTX2 (A) and AQP1 (B) are shown at days P7 in the CP in *Mcidas*$^{-/-}$ and wild type animals. Scale bars, 50 µm. (C) RNAscope analysis of *Gemc1* and *Foxj1* expression is shown at day P7 in the CP in *Mcidas*$^{-/-}$ (black arrowheads) and wild type (black arrows) animals. Ependymal cells lining the ventricles are shown in *Gemc1*$^{-/-}$ (red arrowheads) and wild type (red arrows) animals. Scale bar, 50 µm. (D) The expression of TAp73 is shown at day P7 in the CP in *Mcidas*$^{-/-}$ (black arrowheads) and wild type (black arrows) animals. Ependymal cells lining the ventricles (marked by dotted lines) are shown in *Gemc1*$^{-/-}$ (red arrowheads) and wild type (red arrows) animals. Scale bar, 50 µm.

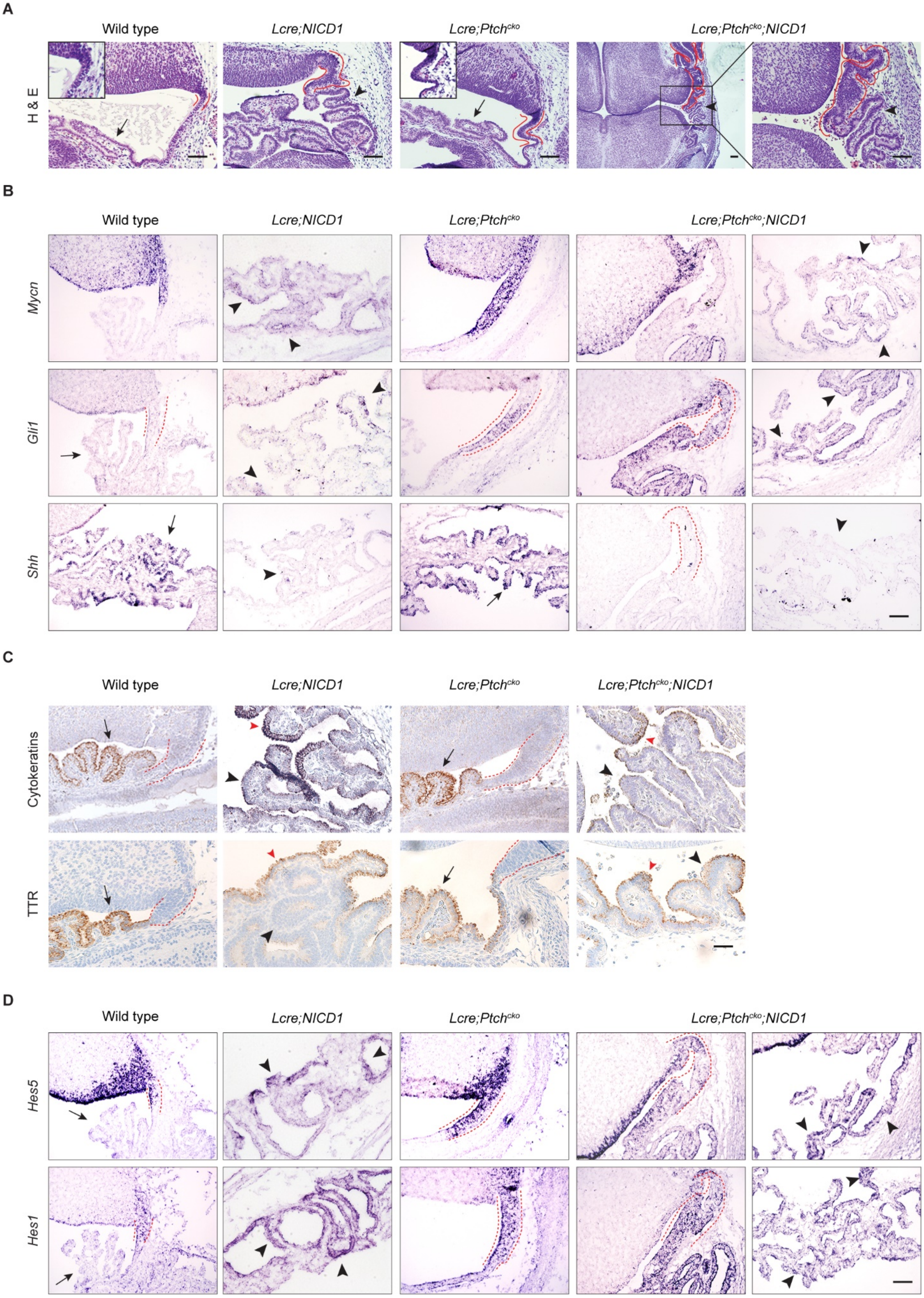


**Supplemental Figure S5**

**Supplemental Figure S5. Analysis of gene expression in CPC driven by aberrant NOTCH and SHH signaling.** (A) H&E staining of coronal sections of hindbrain roof plate/CP (arrows) is shown at day E14.5 in wild type and *Lcre;Ptch^cko^* animals, and abnormal CP growth (arrowheads) in *Lcre;NICD1*, and *Lcre;Ptch^cko^;NICD1* mice. Red lines mark the roof plate magnified in inset images. Boxed region of roof plate/CP in a *Lcre;Ptch^cko^;NICD1* animal is shown in higher magnification on the right. Scale bars,100 µm. (B, C) *In situ* hybridization of *Gli1*, *Mycn* and *Shh* mRNAs (B), or immunohistochemical analysis of the expression of cytokeratins and TTR (C), is shown at day E14.5 in roof plate (marked by dotted lines) and CP (arrows) in wild type and *Lcre;Ptch^cko^* animals, and tumor cells (black arrowheads) in *Lcre;NICD1*, and *Lcre;Ptch^cko^;NICD1* animals. Scale bars, 50 µm. Cytokeratins-expressing and $TTR^+$ epithelial cells (red arrowheads) are mixed with tumor cells in *Lcre;NICD1* and *Lcre;Ptch^cko^;NICD1* animals. Scale bars, 50 µm. (D) *In situ* hybridization analysis of *Hes1* and *Hes5* expression is shown at day E14.5 in hindbrain roof plate (marked by dotted lines) and CP (arrows) in wild type and *Lcre;Ptch^cko^* animals, and tumor cells (arrowheads) in *Lcre;NICD1*, and *Lcre;Ptch^cko^;NICD1* animals. Scale bar, 50 µm.

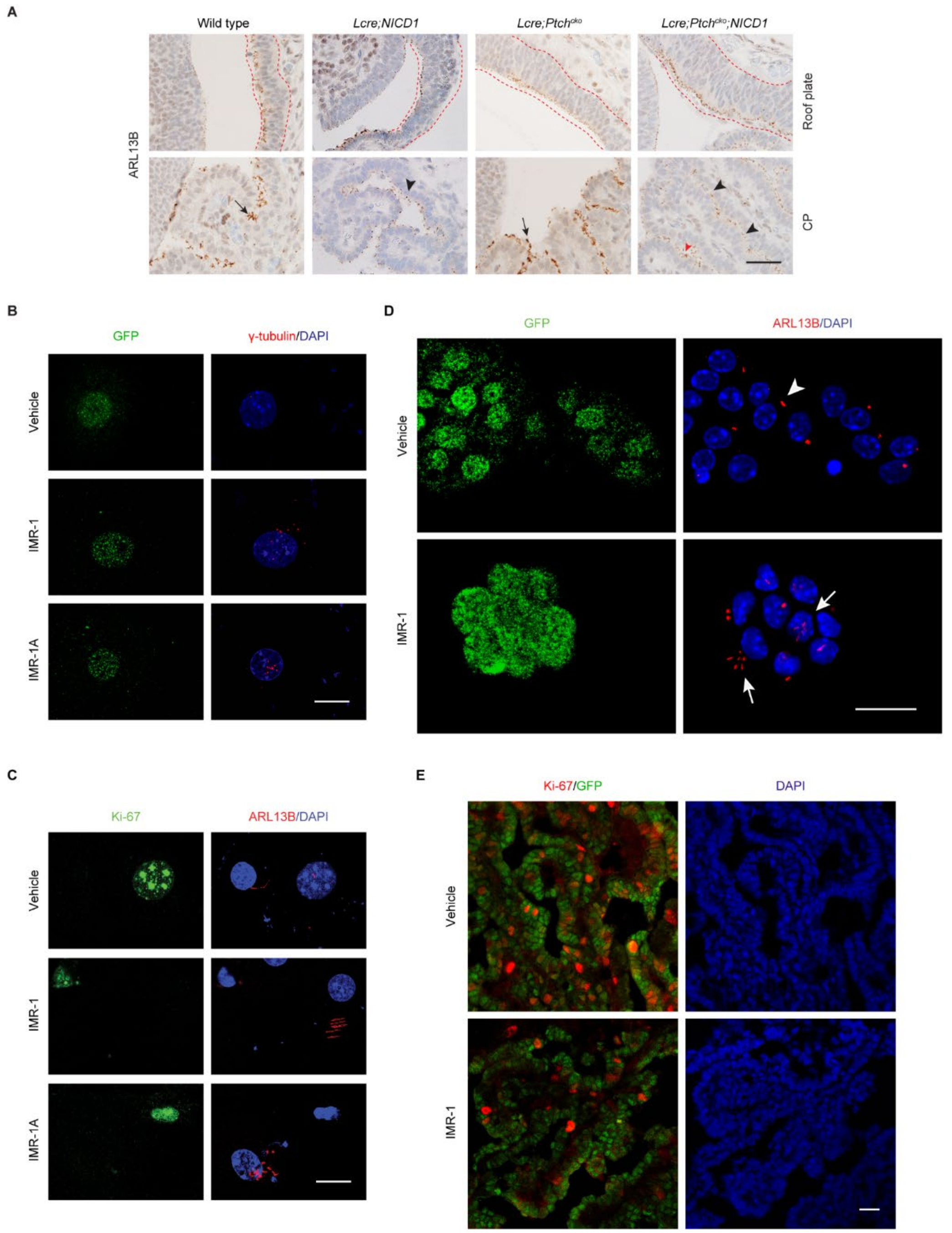


**Supplemental Figure S6. NOTCH inhibition restores multiciliation in CP tumors.** (A) Immunohistochemical analysis of ARL13B expression is shown at day E14.5 in roof plate (marked by dotted lines, top panels) and CP epithelial cells (arrows, bottom panels) in wild type and *Lcre;Ptch$^{cko}$* animals, and tumor cells (black arrowheads, bottom panels) in *Lcre;NICD1* and *Lcre;Ptch$^{cko}$;NICD1* animals. Multiciliated CP epithelial cells (red arrowhead, bottom right panel) are mixed with tumor cells in *Lcre;Ptch$^{cko}$;NICD1* animals. Scale bar, 50 µm. (B, C) The expression of γ-tubulin (B, red), Ki-67 (C, green), and ARL13B (C, red) is shown in tumor cells treated with IMR-1/IMR-1A, or vehicle. GFP (B, green) labels tumor cells, DAPI staining (blue) labels nuclei. Scale bars, 20 µm. (D) *Lcre;NICD1* animals were treated with vehicle or IMR-1 from day E10.5 for 7 days followed by dissociation of tumor cells at day P7. The expression of ARL13B (red) is shown in NICD1$^{+}$/GFP$^{+}$ (green) tumor cells. Primary cilia of tumor cells treated with vehicle (arrowhead) or IMR-1 (arrows) are shown. DAPI staining (blue) labels nuclei. Scale bar, 20 µm. (E) The expression of Ki-67$^{+}$ (red) is shown at day E17.5 in NICD1$^{+}$/GFP$^{+}$ (green) tumor cells from *Lcre;NICD1* animals treated with vehicle or IMR-1 from day E10.5 for 7 days. DAPI staining (blue) labels nuclei. Scale bar, 20 µm.

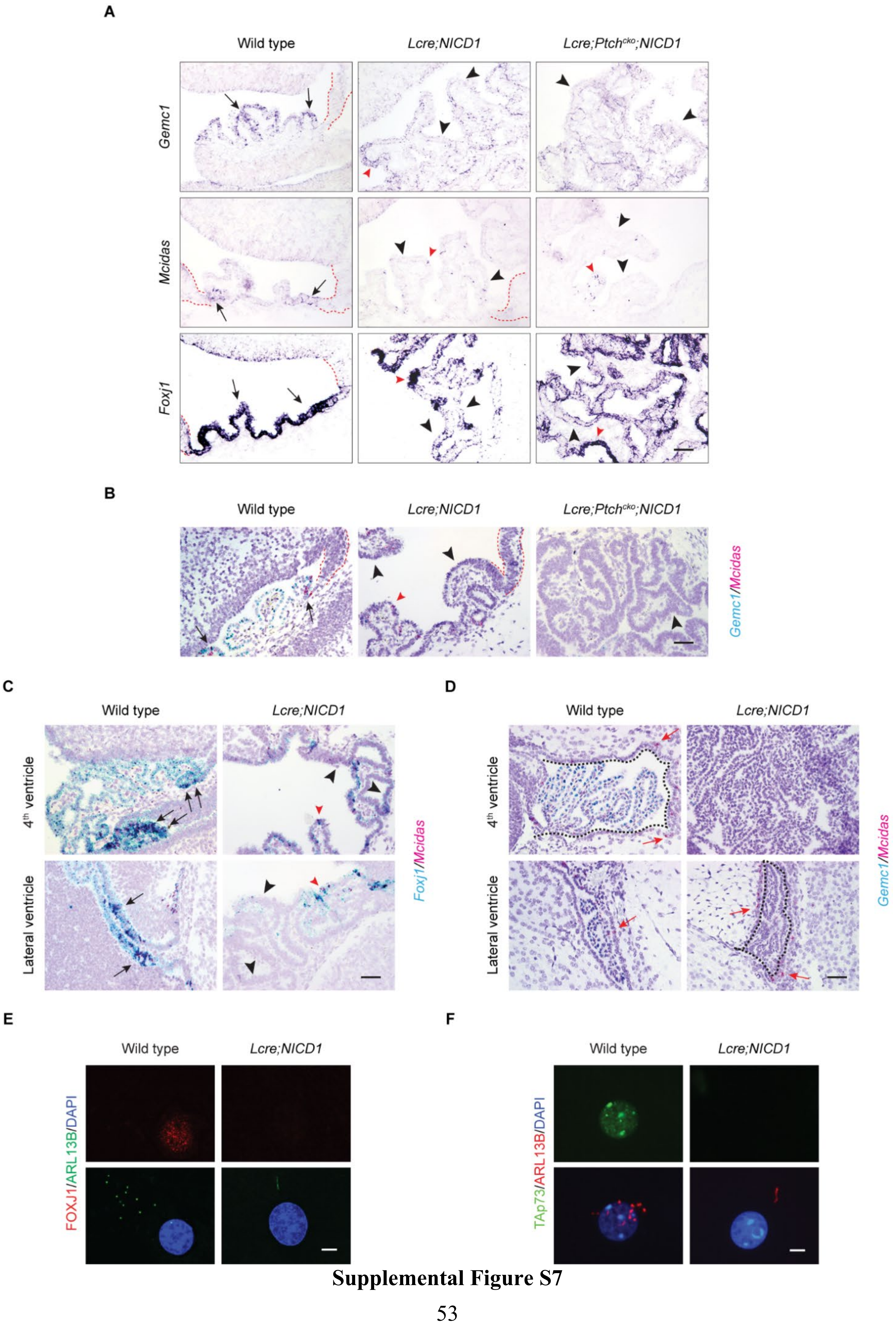


**Supplemental Figure S7**

**Supplemental Figure S7. Repression of *Gemc1-Mcidas* activity in NOTCH-driven CP tumor.** (A) *In situ* hybridization *Gemc1*, *Mcidas*, and *Foxj1* mRNAs is shown at day E14.5 in the roof plate (marked by dotted lines) and CP (arrows) of wild type animals, and tumor cells (black arrowheads) in *Lcre;NICD1* and *Lcre;Ptch$^{cko}$;NICD1* animals. *Gemc1*$^{+}$, *Mcidas*$^{+}$, or *Foxj1*$^{+}$ epithelial cells (red arrowheads) are mixed with tumor cells. Scale bar, 50 µm. (B) RNAscope analysis of *Gemc1* and *Mcidas* expression is shown at day E14.5 in the roof plate (marked by dotted lines) and CP (arrows) in wild type animals, and tumor cells (black arrowheads) in *Lcre;NICD1* and *Lcre;Ptch$^{cko}$;NICD1* animals. *Gemc1*$^{+}$/*Mcidas*$^{+}$ epithelial cells (red arrowheads) are mixed with tumor cells. Scale bar, 50 µm. (C) RNAscope analysis of *Mcidas* and *Foxj1* expression is shown at day E14.5 in roof plate and CP (arrows) in wild type animals, and tumor cells (black arrowheads) in *Lcre;NICD1* animals. *Mcidas*$^{+}$/*Foxj1*$^{+}$epithelial cells (red arrowheads) are mixed with tumor cells. Scale bar, 50 µm. (D) RNAscope analysis of the expression of *Gemc1* and *Mcidas* is shown at day P7 in CP in wild type animals, and tumor cells in *Lcre;NICD1* animals. Scale bar, 50 µm. Ependymal cells in the walls lining the ventricles (marked by dotted lines) are shown that *Mcidas*-positive (red arrows). Scale bar, 50 µm. (E, F) The expression of FOXJ1 (E, red), TAp73 (F, green), and ARL13B (E, green; F, red) is shown in cultured wild type CP epithelial cells and tumor cells from *Lcre;NICD1* animals. DAPI staining (blue) labels nuclei. Scale bars, 5 µm.

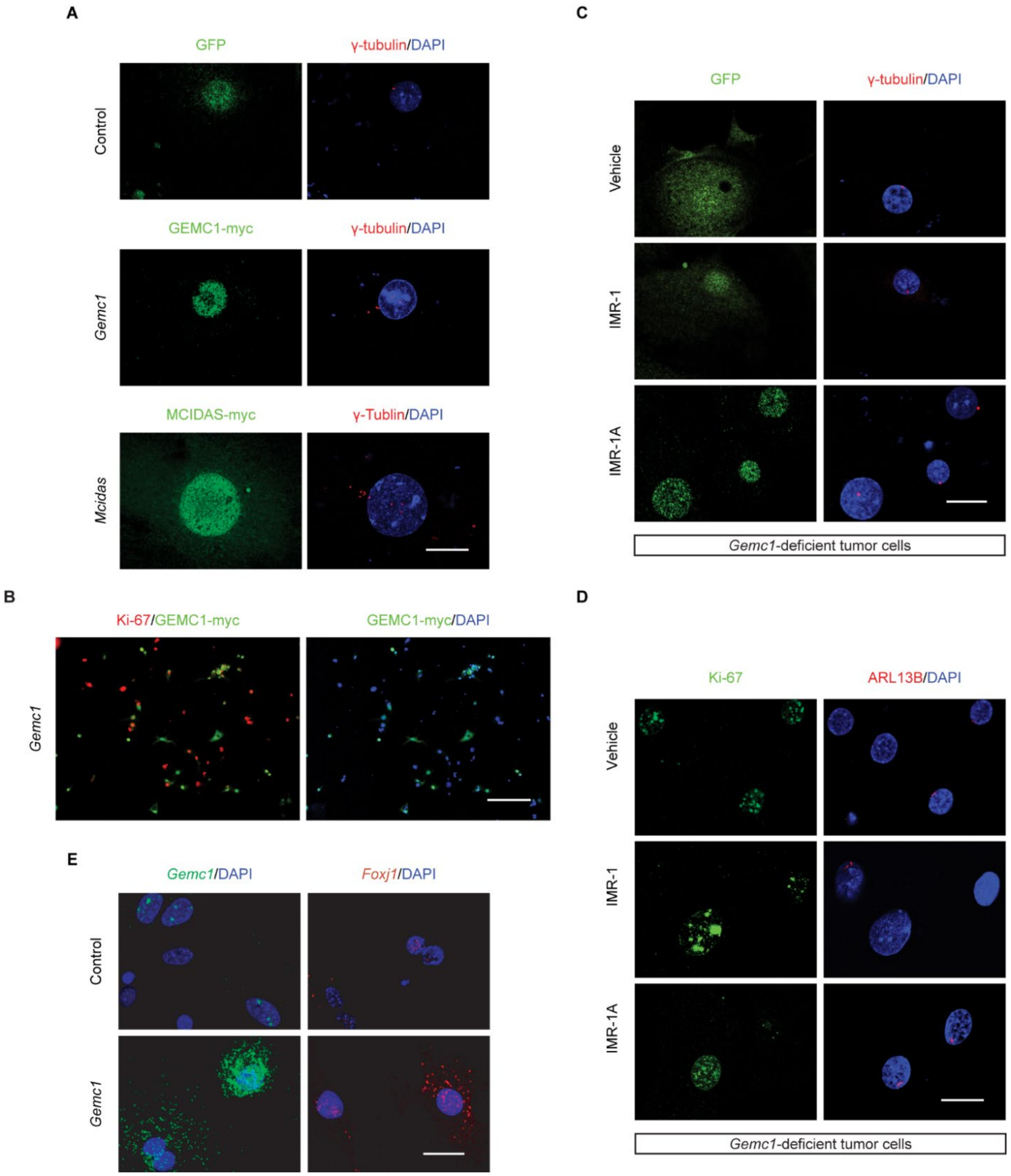


**Supplemental Figure S8.** ***Gemc1*** **suppression mediates multiciliation defects in NOTCH-driven CP tumor.** (A) The expression of γ-tubulin (red) is shown in tumor cells infected with viruses expressing GEMC1-myc or MCIDAS-myc. GEMC1-myc (green), MCIDAS-myc (green), or GFP (green) labels infected or control tumor cells. DAPI staining (blue) labels nuclei. Scale bar, 20 µm. (B) The expression of Ki-67 (red) is shown in tumor cells infected with viruses expressing GEMC1-myc. GEMC1-myc (green) or GFP (green) labels infected or control tumor cells, respectively. DAPI staining (blue) labels nuclei. Scale bar, 20 µm. (C, D) The expression of γ-tubulin (C, red), Ki-67 (D, green), and ARL13B (E, red) is shown in *Gemc1*-deficient tumor cells treated with IMR-1/IMR-1A, or vehicle. GFP (C, green) labels tumor cells. DAPI staining (blue) labels nuclei. Scale bars, 20 µm. (E) RNAscope analysis of *Gemc1* and *Foxj1* expression is shown in *Gemc1*-deficient tumor cells infected with viruses expressing GEMC1-myc or GFP only. DAPI staining (blue) labels nuclei. Scale bar, 20 µm.

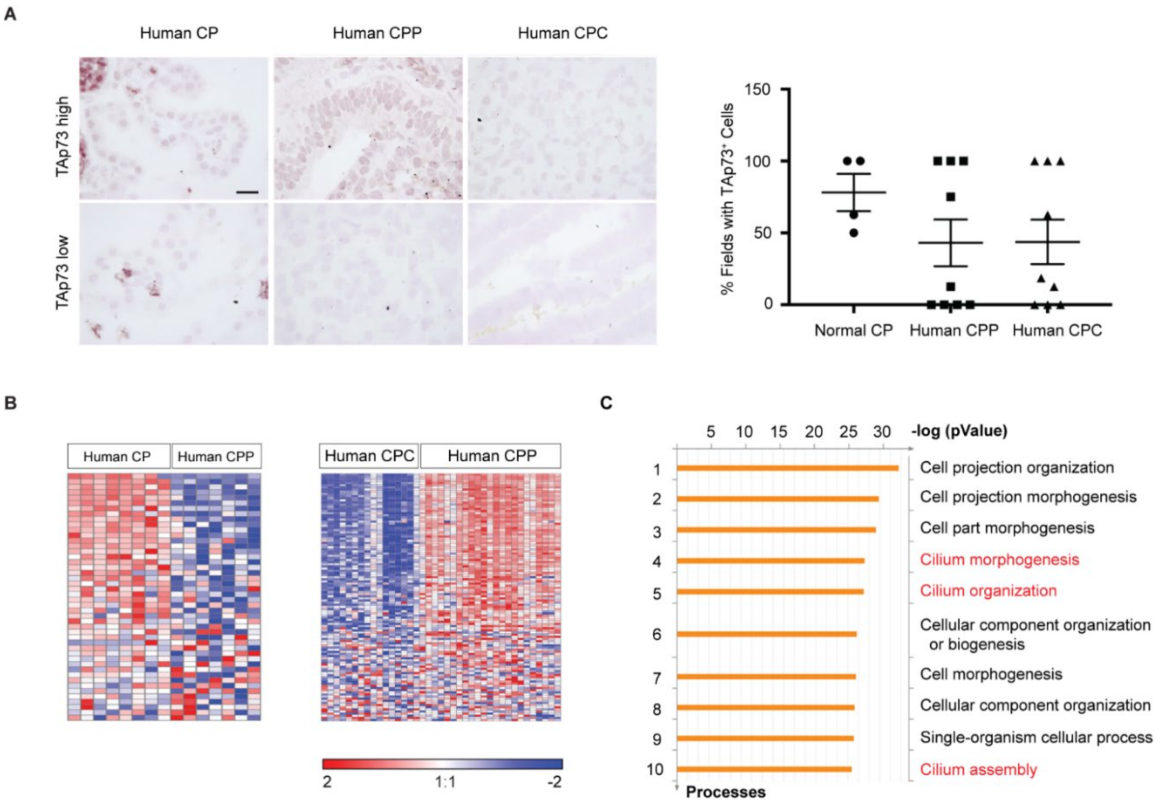


**Supplemental Figure S9. Analysis of gene expression in CP tumors in humans.** (A) Human CP epithelial cells express TAp73, with 50-100% of high magnification fields (100 ×) imaged presenting with TAp73 expression. Human CP tumors displayed variable TAp73 expressions. About half of all CP tumors exhibited TAp73 expression in > 50% of tumor cell population (TAp73 high); the remaining CP tumors display TAp73 expression in small subpopulation of tumor cells (TAp73 low). No difference is observed in TAp73 expression between CPP and CPC. Each point represents one individual and is an average of 4 high-magnification frames across the tumor. Scale bar, 20 μm. (B) Left: hierarchical clustering of human CPPs and normal CPs based on 46 genes involved in cilia differentiation (CPP: $n = 7$ tumors from 7 individuals; normal CP: $n = 8$ CPs from 8 individuals; one-way ANOVA, FDR < 0.05, fold change is shown); right: hierarchical clustering of human CPCs and CPPs with 115 genes involved in ciliogenesis (CPP: $n = 24$ tumors from 24 individuals; CPC: $n = 15$ tumors from 15 individuals; one-way ANOVA, FDR < 0.05, fold change is shown). (C) MetaCore gene enrichment analysis of differentially expressed genes between CPPs and CPCs in humans (CPP: $n = 8$ tumors from 8 individuals; CPC: $n = 15$ tumors from 15 individuals). Significantly enriched signaling networks including cilium morphogenesis, organization, and assembly pathways (red) are shown.

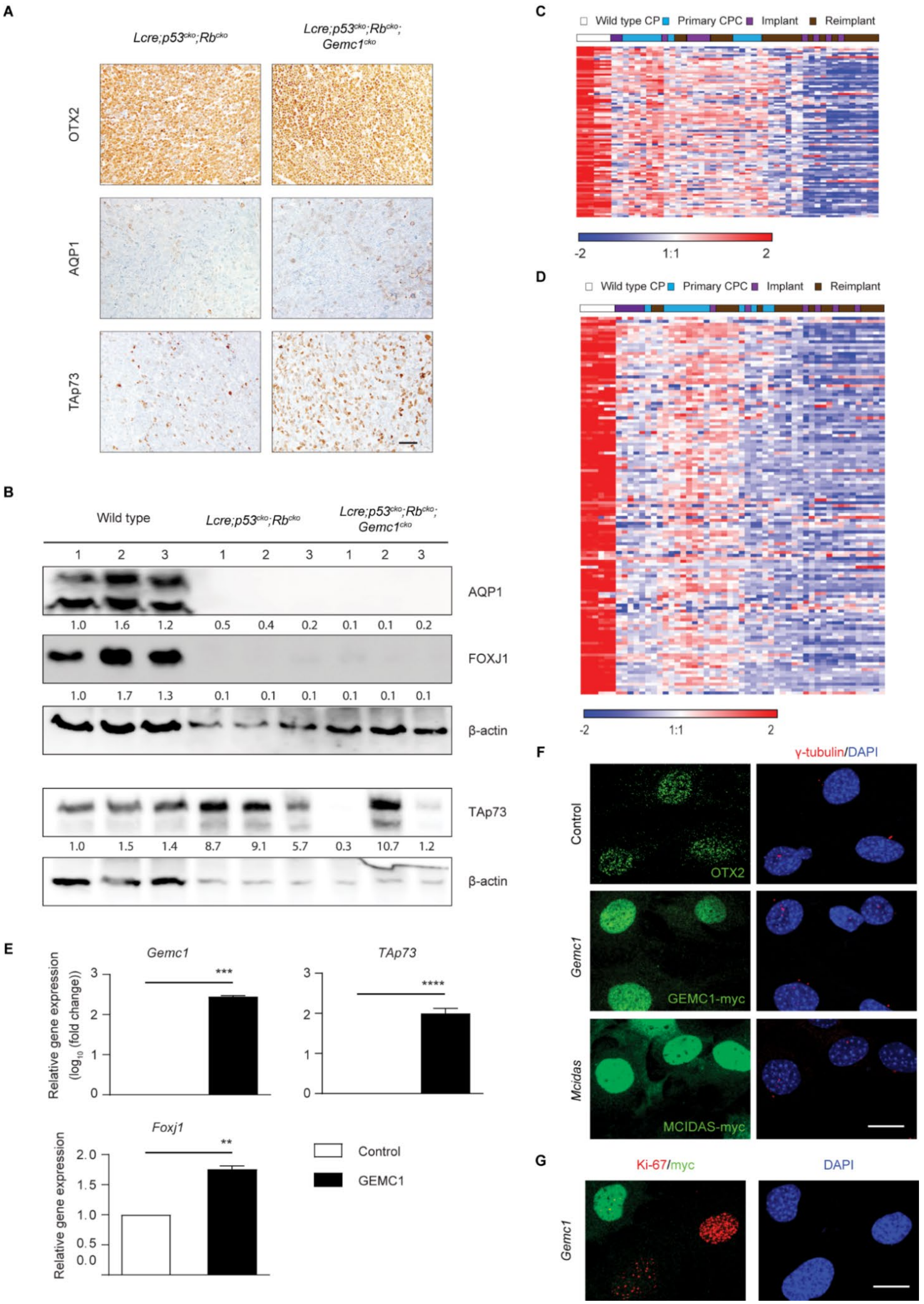


**Supplemental Figure S10**

**Supplemental Figure S10. Analysis of gene expression in *Trp53*-deficient CP tumors.** (A) The expression of OTX2, AQP1 and TAp73 is shown in CP from wild type animals, and tumor cells from *Lcre;p53$^{cko}$;Rb$^{cko}$* and *Lcre;p53$^{cko}$;Rb$^{cko}$*;*Gemc1$^{cko}$* animals. Scale bar, 50 µm. (B) Immunoblot analysis of the expression of AQP1, FOXJ1 and TAp73 in CP from wild type mice, and tumor cells from *Lcre;p53$^{cko}$;Rb$^{cko}$* and *Lcre;p53$^{cko}$;Rb$^{cko}$*;*Gemc1$^{cko}$* mice. The value of each band indicates relative expression level normalized by internal control. (C, D) Hierarchical clustering of CPC driven by *Trp53/Rb1* loss and normal CP based on 88 genes expressed in adult CP epithelial cells (C) or 129 genes involved in ciliogenesis (D) ($n$ = 3 for adult CP, $n$ = 13 for primary tumor, $n$ = 9 for implant, $n$ = 22 for reimplant; one-way ANOVA, FDR < 0.05, fold change is shown). (E) RT-qPCR analysis of tumor cells from *Lcre;p53$^{cko}$;Rb$^{cko}$*;*Gemc1$^{cko}$* animals infected with control viruses or viruses expressing GEMC1-myc ($n$ = 3 samples per treatment, mean ± s.e.m., two-tailed unpaired *t*-test, **$P < 0.01$; ****$P < 0.0001$). (F, G) The expression of ARL13B (F, red) and Ki-67 (G, red) is shown in tumor cells from *Lcre;p53$^{cko}$;Rb$^{cko}$*;*Gemc1$^{cko}$* infected with viruses expressing GEMC1-myc or MCIDAS-myc. OTX2 (green) or GEMC1-myc (green) labels tumor cells. DAPI staining (blue) labels nuclei. Scale bars, 20 µm.
